# Accurate plasmid reconstruction from metagenomics data using assembly-alignment graphs and contrastive learning

**DOI:** 10.1101/2025.02.26.640269

**Authors:** Pau Piera Líndez, Lasse Schnell Danielsen, Iva Kovačić, Marc Pielies Avellí, Joseph Nesme, Lars Juhl Jensen, Jakob Nybo Nissen, Søren Johannes Sørensen, Simon Rasmussen

## Abstract

Plasmids are extrachromosomal DNA molecules that enable horizontal gene transfer in bacteria, often conferring advantages such as antibiotic resistance. Despite their significance, plasmids are underrepresented in genomic databases due to challenges in assembling them, caused by mosaicism and micro-diversity. Current plasmid assemblers rely on detecting circular paths in single-sample assembly graphs, but face limitations due to graph fragmentation and entanglement, and low coverage. We introduce PlasMAAG (Plasmid and organism Metagenomic binning using Assembly Alignment Graphs), a framework to recover plasmids and organisms from metagenomic samples that leverages an approach that we call “assembly-alignment graphs” alongside common binning features. On synthetic benchmark datasets, PlasMAAG reconstructed 50–121% more near-complete plasmids than competing methods and improved the Matthews Correlation Coefficient of geNomad contig classification by 28–106%. On hospital sewage samples, PlasMAAG outperformed all other methods, reconstructing 33% more plasmid sequences. PlasMAAG enables the study of organism-plasmid associations and intra-plasmid diversity across samples, offering state-of-the-art plasmid reconstruction with reduced computational costs.

## INTRODUCTION

Plasmids are extrachromosomal DNA molecules within a host cell that are physically separated from chromosomal DNA and can replicate independently (1–3). Plasmids differ in genome length, copy number, replication mechanism, and conjugation mode. The part of the plasmid that encodes the core replication machinery is typically contiguous and is called the ‘backbone’. The replication and maintenance of plasmids incur a metabolic burden for the host, and to avoid purifying selection, the plasmid must carry additional ‘cargo’ genes that increase the fitness of either the host or the plasmid (4). This may be genes that confer antibiotic resistance (5–7).

Approximately 50% of bacteria carry one or more plasmids (8). Nonetheless, in databases, sequences from plasmids remain underrepresented compared to those from cellular genomes. For instance, RefSeq contains 82,471 bacterial genomes, but only 7,892 plasmids (9). Characterization of environmental plasmids in *in vitro* conditions is inherently limited by the so called “cultivation bottleneck” (10, 11), where laboratory conditions modify microbial diversity, offering a poor representation of the original composition. Therefore, despite the great number of plasmid-related studies, most studies have investigated plasmid virulence and properties in isolated strains (8, 12–14). This mismatch between estimated plasmid prevalence in bacteria and plasmid representation in the databases emphasizes our incomplete understanding of the plasmid genetic structure, diversity, and function (2).

Metagenomic offers culture free alternative techniques; however, the genetic complexity of environmental samples complicates the process (10, 11, 15). Besides the challenges of assembling bacterial chromosomes, assembly of plasmids bring additional challenges: a) plasmids undergo frequent recombination, creating groups of plasmids that share a ‘backbone’ but diverge on their ‘cargo’ sequence (12); b) plasmids at high copy number have higher mutation rates than chromosomes, which increases micro-diversity and makes them difficult to assemble with de Bruijn-graph based assemblers (16) and c) plasmids are enriched for repeated sequences associated with transposable elements (2). A consequence of this is that plasmid sequences will be fragmented across assemblies and entangled by sharing the same ‘backbone’ and repeated genetic elements (12, 17).

To overcome these challenges, dedicated metagenomic plasmid assemblers such as Recycler, metaplasmidSPAdes, and SCAPP have been developed (18–20). These methods rely on the assembly graph, a data structure used by metagenome assemblers, that represents overlaps between sequencing reads. Assembly graphs represent contiguous sequences (contigs) as nodes and overlaps between these as edges (21). By leveraging assembly graphs, plasmid assemblers can identify connected sequences and resolve complex genomic regions (22). Recycler re-interprets the metagenomic assembly graph, leveraging paired-reads information, and attempting to extract subgraph cycles with uniform coverage from the graph, in a process named graph ‘peeling’ also used by SCAPP (18). MetaplasmidSPAdes iteratively extracts cyclic subgraphs from the assembly graph with uniform coverage, filtering the subgraphs with plasmidVerify (19), a tool that classifies sequences into plasmidic and chromosomal based on gene content using a profile-HMM (19). Finally, SCAPP tries to find plasmid cyclic paths in assembly graphs based on paired read mappings, presence of plasmid-specific genes, sequence length, coverage, and plasmid sequence score annotation based on PlasClass (18, 23). Common to the methods is that they operate on single-sample assembly graphs, use the circularity of plasmids, and contig coverage. However, the methods have fundamental limitations. First, low coverage causes the “fragmentation problem”, where some plasmids appear disconnected in the graph, making them impossible to identify by graph peeling (14, 24). Second, the high recombination rate of plasmids causes entangled assembly graph components where circularity is hard to identify (25). Finally, SCAPP and metaplasmidSPAdes leverage plasmid gene signatures to guide the plasmid candidate’s path extraction from the assembly graph.

Binning is a computational strategy used to reconstruct genomes by grouping contigs based on their genome of origin, providing an alternative to assembly graph-based methods. Modern binners typically integrate several sequence features, including k-mer composition (26–31), abundance patterns across samples (26–31), assembly graph connectivity (32), and taxonomic markers or annotations (28, 30, 33). Most of these features can be computed on a per-sequence basis and are therefore not vulnerable to the fragmentation problem suffered by assembly graphs with low coverage. Furthermore, it has been shown how binning information can be used to refine contig classification, using binning features to guide classification rather than contig classification to reconstruct the original sequences (34, 35). We have previously developed the binning tool VAMB, which combines several of these features using a variational autoencoder into a latent space which is clustered to form bins (26, 36).

In this paper, we introduce assembly-alignment graphs (AAGs), which combine the intra-sample sequence overlaps recorded by assembly graphs, with cross-sample overlaps detected by ordinary sequence alignment. Using a contrastive learning approach, we were able to effectively integrate the AAG with ordinary binning features in a new binning framework called PlasMAAG that can reconstruct both plasmids and cellular genomes. We evaluated PlasMAAG on simulated data, where it reconstructed 9-70% more near-complete (≥0.95 precision, ≥0.9 recall) (NC) plasmids and cellular genomes than SemiBin2, ComeBin, MetaBAT2, MetaDecoder, and VAMB. Regarding only plasmids, PlasMAAG reconstructed at least 50-121% more NC plasmids than any other binner, and 14-40% more NC plasmids than the unfiltered set of cycles from SCAPP cycles in 4/5 benchmark datasets. When using a confident threshold PlasMAAG reconstructed 21-212% more confident NC plasmids than SCAPP confident. PlasMAAG achieved excellent performance on hospital sewage samples, reconstructing at least 33% more plasmid sequences than any other tool, as evaluated with a robust paired long-read short-read validation setup. Using PlasMAAG’s ability to reconstruct plasmids and hosts, we studied host-plasmid associations in hospital sewage samples and intra-plasmid diversity across samples. To our knowledge, PlasMAAG is the only method that enables ‘multi-sample’ characterization of both plasmids and cellular genomes, achieving state-of-the-art plasmid reconstruction with reduced computational resource requirements than current plasmid binners.

## RESULTS

### PlasMAAG: Combining assembly graphs, alignment graphs, TNFs, and co-abundances for binning

PlasMAAG is a new deep learning binning algorithm designed to reconstruct cellular and plasmid genomes (**Figure 1**). Compared to our previous developed binning algorithm VAMB, PlasMAAG introduces three novelties. First, we combine multi-sample assembly graphs with contig alignment graphs into a single graph called ‘assembly-alignment graph’. The assembly-alignment graph is then projected to an embedding space with fastnode2vec (37), from which communities of contigs can be extracted. Second, we enhanced the training of the variational autoencoder (VAE) by adding contrastive learning, based on information from the assembly-alignment graph. Finally, we leverage binning to ensemble geNomad (38) contig annotation scores across bins to classify the bins into plasmid or cellular genomes.

**Figure 1.**
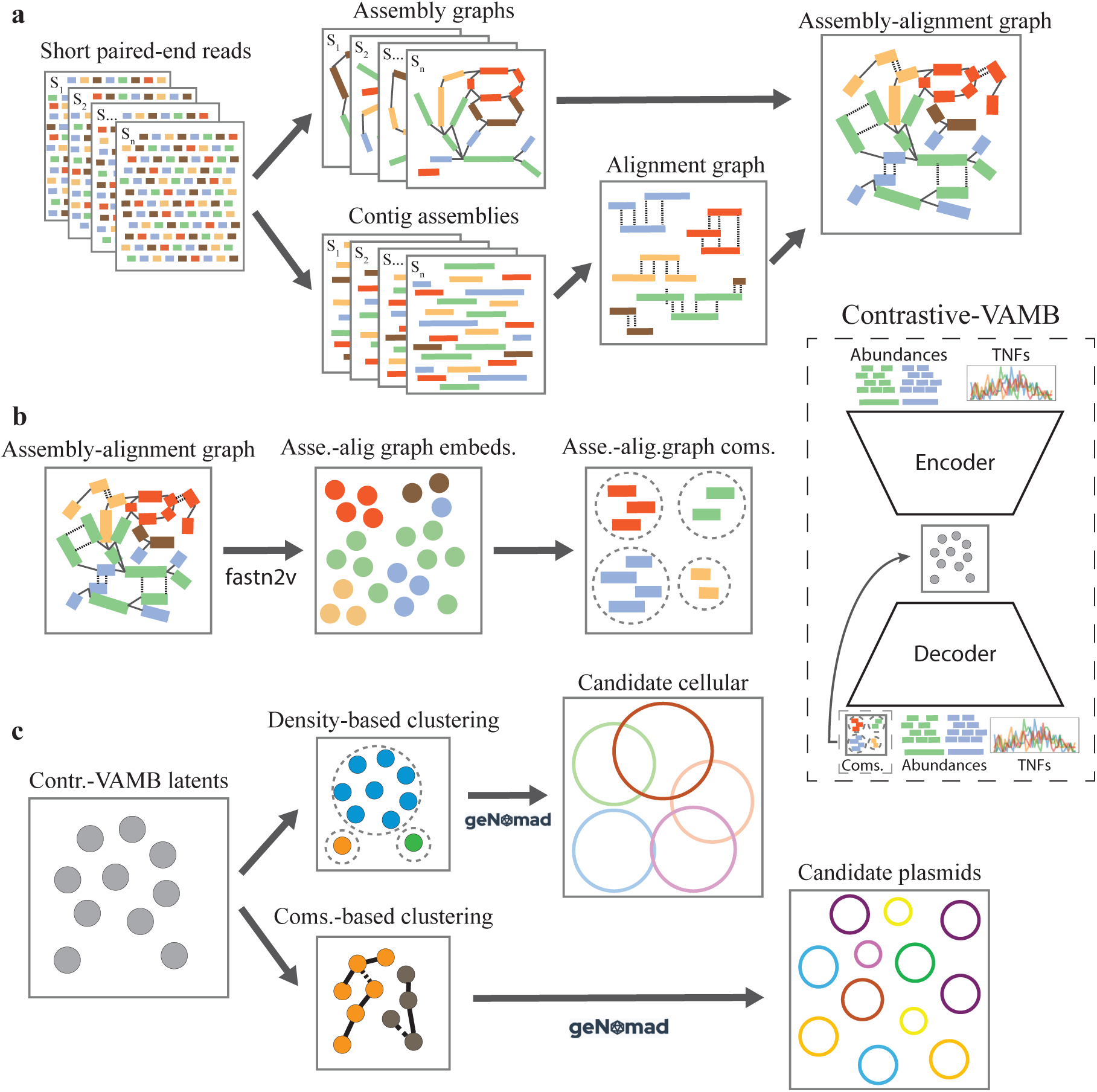
PlasMAAG workflow overview. PlasMAAG leverages assembly graphs, alignment graphs, k-mer signal, and contig co-abundances for binning, with a final step where bins are classified as cellular or plasmid bins based upon refined geNomad predictions. **a**. Per-sample assembly graphs are merged with the between-sample alignment graph, generating the assembly-alignment graph. **b**. Fastnode2vec is used to generate contig embeddings from the assembly-alignment graph, from where contig communities are extracted. Communities are expanded, merged, and purified using a variational autoencoder with contrastive loss that push communities towards be preserved in the embedding. **c**. Plasmid and cellular candidate bins are extracted from the VAE embedding based on their geNomad scores, using distinct plasmid and cellular clustering strategies.

### PlasMAAG reconstructed 21-212% more plasmid bins compared to SCAPP confident

To develop and test PlasMAAG we re-assembled the simulated CAMI2 short-read human microbiome toy datasets. Re-assembly of CAMI2 was required because the original CAMI2 plasmids were not simulated as independent entities from their hosts cellular genomes, and because assembly graphs were not available (see Methods). We found that PlasMAAG reconstructed 5-64% more NC bins over all 5 benchmark datasets compared to VAMB, the second best performing binner on the benchmark data **(Figure 2.A).** The improvement in binning performance was driven by increased reconstruction of plasmids, where PlasMAAG reconstructed at least 50-121% more NC plasmids candidate bins compared to SemiBin2, ComeBin, MetaBAT2, MetaDecoder, and VAMB across all benchmark datasets. When comparing to SCAPP, PlasMAAG reconstructed 14-40% more NC plasmids than SCAPP cycles over 4/5 benchmark datasets **(Figure 2.B)**. When evaluating confident plasmids bins generated by PlasMAAG (above 0.95 geNomad plasmid threshold), PlasMAAG reconstructed 21-212% more NC plasmid bins than SCAPP confident (**Figure 2.C**). Furthermore, PlasMAAG spanned a larger variation of plasmids, since the unique set of confident PlasMAAG plasmids across the benchmark datasets included 172 NC plasmid bins and 223 medium-quality (≥0.9 precision, ≥0.5 recall) (MQ) plasmid bins not reconstructed by SCAPP confident. In contrast, SCAPP confident reconstructed 64 NC plasmid bins and 68 MQ plasmid bins not reconstructed by PlasMAAG. The intersecting set of plasmids reconstructed by both methods was 164 NC, and 185 MQ plasmid bins, respectively (**Figure 2.D-E**). Considering cellular binning, PlasMAAG was also competitive, reconstructing 0.7-9% less NC cellular bins than VAMB, the best cellular binner on benchmark datasets **(Figure 2.F)**. The set of PlasMAAG confident plasmids offered a better balance than SCAPP confident between the true positive and true negative plasmids present in the benchmark datasets, with a 14-43% improvement in F1 (**Figure 2.G-H**, **Supplementary Figure 1**, **Supplementary Note 1**). By averaging geNomad scores across PlasMAAG’s clusters, we can detect plasmids more accurately than applying geNomad on individual contigs, yielding an improvement over the plasmid/non-plasmid contig classification Area Under Precision-Recall Curve (AUPRC) and Matthews correlation coefficient (MCC) of between 28-69% and 42-131%, respectively (**Figure 2.I-J**, **Supplementary Note 2**, **Supplementary Figure 2**, **Supplementary Table 1**).

**Figure 2.**
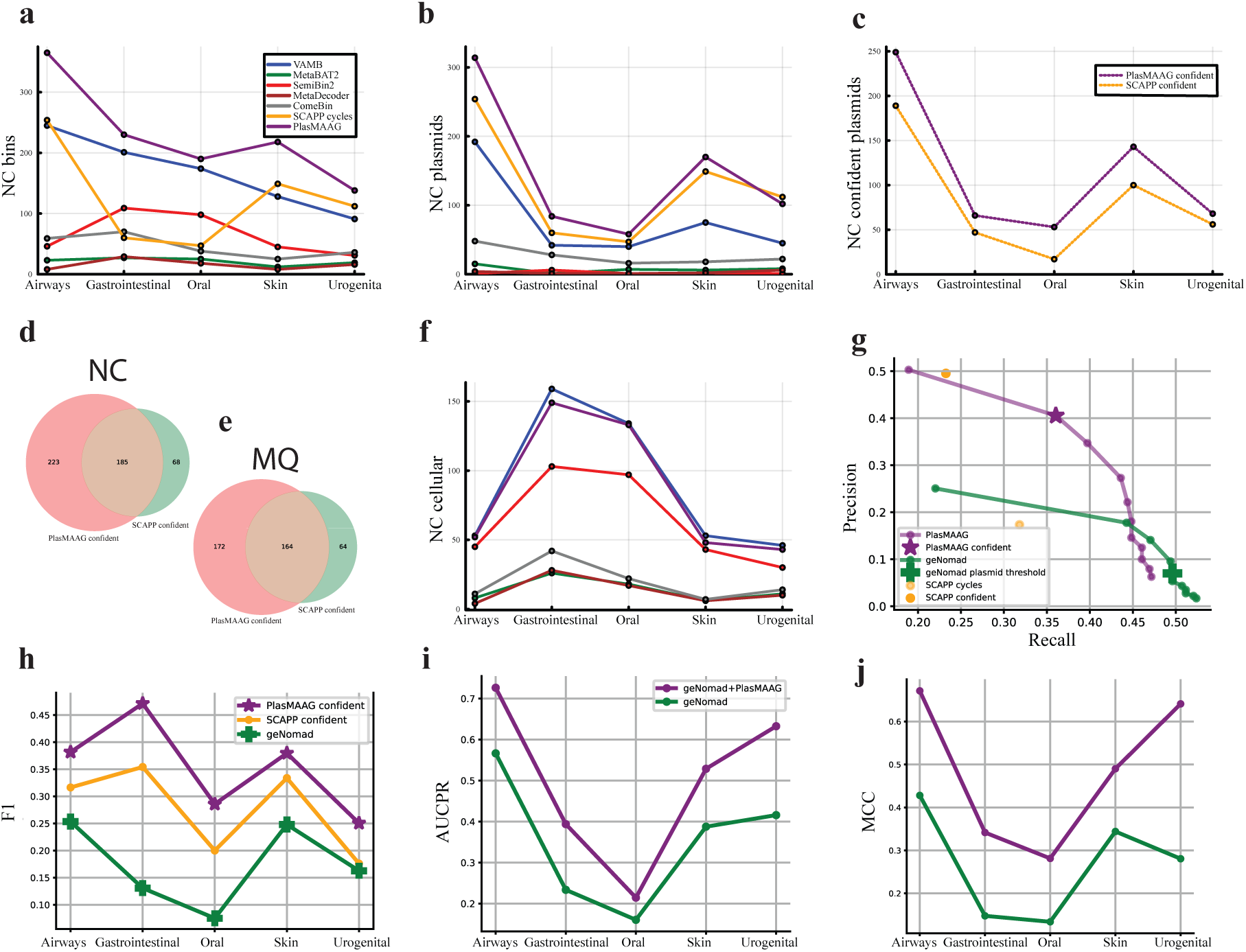
PlasMAAG binning and classification performance across the benchmark datasets. **a.** NC bins (cellular + plasmids) reconstructed from the five benchmark datasets for VAMB (blue), MetaBAT2 (green), SemiBin2 (red), MetaDecoder (brown), ComeBin (grey), SCAPP cycles (yellow), and PlasMAAG (purple). **b**. NC plasmid bins reconstructed by all methods. **c**. NC plasmids reconstructed by SCAPP confident (yellow dotted) and PlasMAAG confident (purple dotted). **d**. Set of NC complete unique plasmids reconstructed only by PlasMAAG confident (red), only by SCAPP confident (green), and by both methods (light brown) across all datasets. **e**. Same than **d** but for MQ plasmid bins. **f**. NC cellular bins reconstructed by all methods except SCAPP confident. **g**. Plasmid sample precision-recall (see Methods) from the Airways dataset for PlasMAAG across geNomad thresholds (purple), PlasMAAG confident (purple star), geNomad across thresholds (green), geNomad at the default plasmid threshold (green cross), SCAPP cycles (light yellow), and SCAPP confident (dark yellow). **h**. Sample F1 across the five benchmark datasets for geNomad at the default plasmid threshold (green), SCAPP confident (yellow), and PlasMAAG confident (purple). **i**. Area Under Precision-Recall Curve (AUPRC) for the classification of plasmids by geNomad (green) and when aggregating the geNomad scores per PlasMAAG community-based clusters (purple). **j**. Matthew correlation coefficient (MCC) for the classification of plasmids by geNomad (green) and when aggregating the geNomad scores per PlasMAAG community-based clusters (purple).

### Assembly graphs have a strong signal for binning

In assembly graphs, edges represent sequence overlaps between contigs. Therefore, it has long been known that they are informative for binning (32, 39). To quantify how informative edges were, we weighted them by *normalized linkage* (see Methods), based on the number of overlapping k-mers, and the length of the contigs. Normalized linkage showed a positive correlation with edge accuracy at genome (species) level, with Spearman correlation coefficients 0.49-0.93 (0.86-0.98) across all benchmark datasets (**Figure 3.A**, **Supplementary Figure 3**). Additionally, normalized linkage was evaluated for correlation with edge accuracy (i.e. how often two contigs linked by an edge belong to the same genome) by calculating the Area Under Precision-Recall Curve (AUPRC). The resulting AUPRC ranged from 0.66 to 0.74 at the genome level and 0.81 to 0.90 at the species level across the benchmark datasets (**Figure 3.B**, **Supplementary Figure 3**). We concluded that the assembly graph contains useful signals for binning.

**Figure 3.**
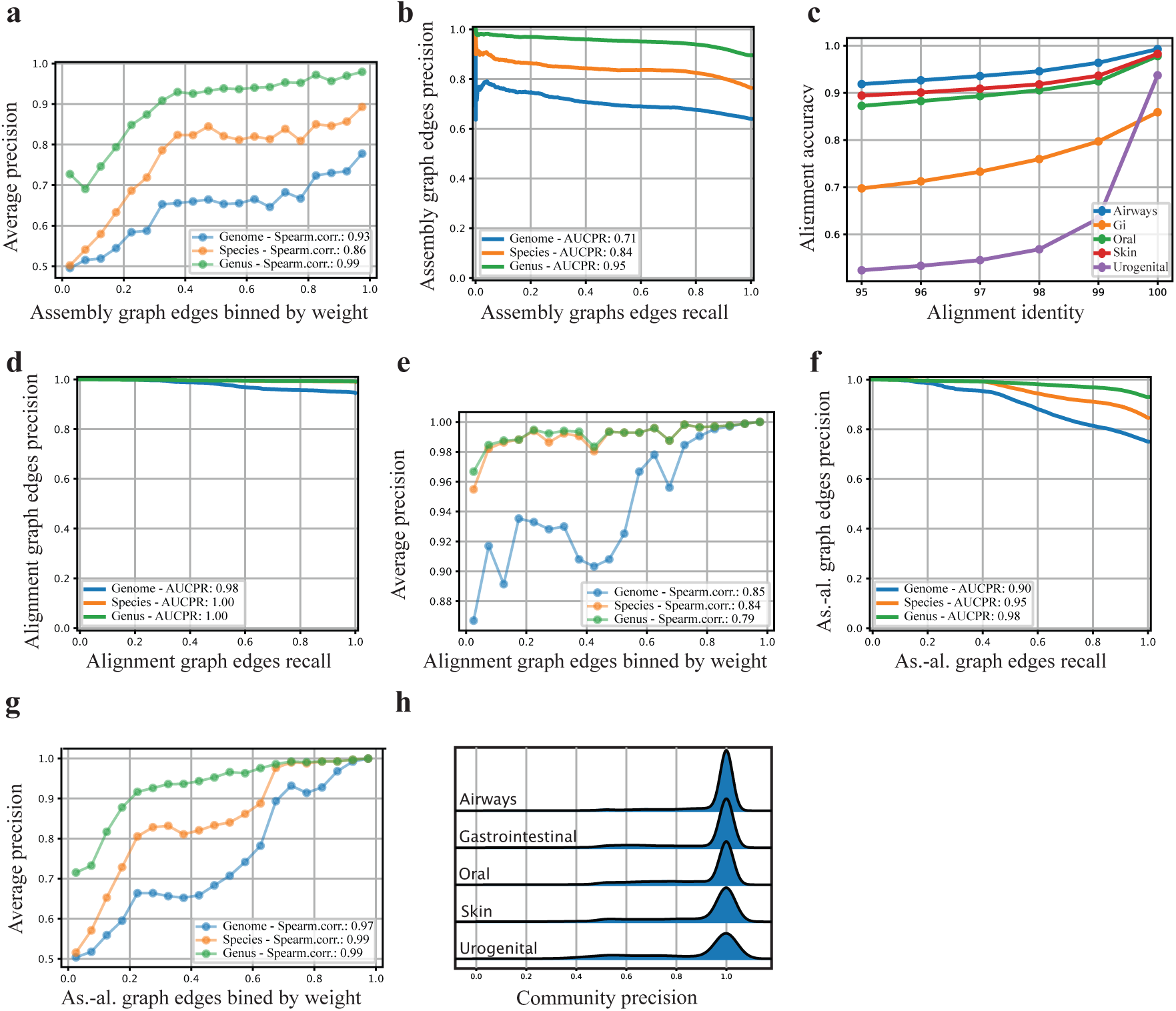
Assembly graph, alignment graph, and assembly-alignment graph-based features for binning. **a**. Average precision of the assembly graph edges from the Airways benchmark dataset, sorted by edge weight and grouped into 5% bins, is shown for genome (blue), species (orange), and genus (green) taxonomic levels. **b**. Precision-recall curve of the assembly graph edge weights from the Airways benchmark dataset at genome (blue), species (orange), and genus (green) taxonomic levels. **c**. Alignment accuracy when increasing minimum identity thresholds across benchmark datasets. Results are shown only for restrictive alignments (see Methods) between contigs from different samples. **d**. Precision-recall curve of the alignment graph edge weights from the Airways benchmark dataset at genome (blue), species (orange), and genus (green) taxonomic levels. **e**. Average precision of the alignment graph edges from the Airways benchmark dataset, sorted by weight and grouped into 5% bins is shown for genome (blue), species (orange), and genus (green) taxonomic levels. **f**. Precision-recall curve of the assembly-alignment graph edge weights from the Airways benchmark dataset at genome (blue), species (orange), and genus (green) taxonomic levels. **g**. Average precision of assembly-alignment graph edges from the Airways benchmark dataset, sorted by weight and grouped into 5% bins is shown for genome (blue), species (orange), and genus (green) taxonomic levels. **h.** Precision distribution of communities extracted using FastNode2Vec from assembly-alignment graphs across the five benchmark datasets at genome taxonomic level.

### Alignment graphs contain taxonomic information across samples

PlasMAAG uses the multi-split binning workflow due to its superior accuracy (26, 36), where samples are assembled individually. Therefore, assembly graphs only inform about overlaps between intra-sample contigs. To also include between-sample contig overlap information, we aligned contigs across samples with strict criteria to accept a hit (see Methods). The alignments were highly precise with an accuracy at genome (species) level of 57-95% (95-99%) (**Figure 3.C**, **Supplementary Figure 4**). By adding alignments between pairs of contigs as edges to the assembly graph, we created an alignment-assembly graph (AAG), where we weighed each edge by either alignment metrics (for alignment edges) and normalized linkage (for assembly graph edges, see Methods). Alignment edge weight between two contigs correlated with taxonomic relatedness of the contig’s genomes, showing an 82-98 (98–100) Area Under Precision-Recall Curve (AUPRC) across the benchmark datasets at genome (species) taxonomic level (**Figure 3.D**, **Supplementary Figure 5**). Furthermore, there was a positive correlation between the averaged alignment graph edges and the average accuracy, with a Spearman correlation coefficient of 0.71-0.95 across all benchmark datasets (**Figure 3.E**, **Supplementary Figure 5**).

### Assembly-alignment graphs integrate alignments and assembly graphs

The complementarity between cross-sample alignments and the intra-sample assembly graph connections in the AAG enabled us to integrate these in a unified graph, resulting in a combined graph that we named ‘assembly-alignment graph’. We evaluated the edges in the assembly-alignment graphs across the benchmark datasets to assess whether higher edge weights correspond to contigs that are taxonomically close, such as those from the same genome. The edge weights in the assembly-alignment graph reflect taxonomic relationship between sequences, consistent with the original assembly and alignment graphs, achieving a AUPRC of 0.69-0.90 (0.93-0.97) across the benchmark datasets at genome (species) taxonomic level (**Figure 3.F**, **Supplementary Figure 6**). Consistently with the AUPRC findings, we found a positive correlation between the averaged edge weights and the average edge accuracy at genome taxonomic level, with 0.20-0.97 Spearman correlation coefficients across benchmark datasets (**Figure 3.G**, **Supplementary Figure 6**). The assembly-alignment graph integrates assembly graphs and alignment information across samples into a unified object, where edge weights reflect taxonomic relationships.

### Extracting high precision, low completeness communities from the assembly-alignment graph

The majority of contigs are too short to contain a stable signal for binning, but the AAG cohesion depends on the nodes representing short contigs. Therefore, we condensed the AAG into a set of node communities using fastnode2vec (see Methods). We found that the extracted communities from this graph embedding had high purity, with an average precision at genome (species) level of 86-95% (95-97%) across the benchmark datasets, and where 63-84% (85-91%) of communities had a precision at genome (species) level (**Figure 3.H**, **Supplementary Table 2**). However, we observed that communities were composed of rather few contigs, with 85-91% of the communities were composed of 10 or less contigs across the benchmark datasets. Furthermore, microbial genomes were fragmented in, on average, 12.2-32.8 communities, and plasmids somewhat were less fragmented, split between 1.6-2.5 communities on average (**Supplementary Figure 7**, **Supplementary Table 2**). We also noticed that only 31-47% of contigs in the datasets belonged to any community (**Supplementary Table 2**). In conclusion, the communities extracted from the AAG using fastnode2vec were precise, but incomplete and fragmented.

### Contrastive variational autoencoders improve binning through aggregating, merging and splitting communities

To address the fragmentation of AAG communities, we leveraged traditional binning features such as contig k-mer composition and abundances (40). In the VAMB framework, these contig features are embedded using a variational autoencoder (VAE), and these embeddings are then used to cluster contigs together. PlasMAAG follows the same approach but also considers community structure during the embedding and clustering process. To encourage contigs of the same community to be close in the embedding, we added an extra term to the loss function of the VAE which penalized high embedding distance between contigs of the same community. We call this term ‘contrastive loss’. We then applied a clustering strategy on the contrastive VAMB embeddings, consisting of three key steps: (1) Merging – Communities close in the embedding were merged to reduce genome fragmentation, and increase genome recall. (2) Splitting – Communities with contigs placed far apart in the embedding were split up to increase precision. (3) Expansion – Unassigned contigs located close to a community in latent space were added to the community to improve recall. We refer to these three steps as ‘community-based’ clustering (see Methods, **Supplementary Figure 8**). This community-based clustering resulted in a 46–102% increase in genome recall across benchmark datasets compared to the raw communities, confirming the effectiveness of the community merging step. The splitting step improved precision by 0.03–1% (**Supplementary Figure 9**, **Table 3**), indicating minor but positive impact without compromising recall. On the other hand, community expansion had limited effect, with only a 1–3% increase in community size (**Supplementary Table 3**), suggesting that step 3 had a smaller impact. Since recall increased and precision slightly improved, F1 scores also increased, along with the number of reconstructed near-complete (NC) bins.

### Contrastive loss had a positive impact on binning

To better understand the importance of the contrastive loss on the latent representations, we evaluated how it impacted community-based clustering and clustering from the original VAMB, which we call ‘density-based’ clustering. Community-based clustering with contrastive loss achieved 28–63% higher average F1 scores compared to clustering without the contrastive loss, reconstructing 7–45% more NC bins across the benchmark datasets (**Supplementary Figures 10–11**). Contrastive loss also improved density-based clustering, causing a 57–162% increase in F1 scores across all benchmark datasets (**Supplementary Figure 10**), but did not uniformly increase the number of NC bins. NC bin recovery was increased by 1–6% in 4 out of 5 datasets but led to 16% fewer NC bins in one dataset due to a small decrease in precision (**Supplementary Figure 11**). Overall, the contrastive loss boosted recall and led to significantly higher F1 scores in both clustering approaches, whereas its effect on precision and final NC bin counts varied depending on the dataset and clustering strategy, highlighting the trade-offs introduced by enforcing graph-based community structures in the latent space.

### Differential embeddings of plasmids and organisms requires tailored clustering leveraging geNomad

As previously mentioned, we found the cellular genomes to be fragmented across more communities than plasmids, probably due to the larger size of cellular genomes. Furthermore, we observed distinct patterns in k-mer composition, contig co-abundance, and PlasMAAG latent representations between plasmids and cellular genomes (**Supplementary Note 3**, **Supplementary Figures 12-14**, **Supplementary Tables 3-4**). This suggested that community-based clustering might be more suitable for plasmids, and density-based clustering for cellular genomes, which we indeed verified with our benchmark datasets (**Supplementary Note 3**, **Supplementary Figures 12-14**, **Supplementary Tables 3-4**). Therefore, to identify potential plasmid communities in the AAG, we used geNomad to assign plasmid scores to each community (38). We found that averaging geNomad scores across communities led to more accurate plasmid identification compared to scoring individual contigs (**Supplementary Note 2**, **Supplementary Figure 2**, **Supplementary Table 1**). This allowed us to extract communities as putative plasmid bins for community-based clustering and clustered the remaining contigs using density-based clustering. Additionally, we found that this was sensitive to the geNomad threshold used for the classification, particularly in the case of organisms (**Supplementary Note 4**, **Supplementary Figures 15-16**). For instance, when setting a geNomad plasmid threshold of 0.7, we observed a decrease on the NC cellular genomes (plasmids) of 6-39% (3-18%) (**Supplementary Figure 17**). This indicated that the selection and dereplication process, based on geNomad-identified plasmid clusters, led to a trade-off in cellular genomes recovery. We conclude that integrating geNomad sequence predictions with PlasMAAG’s diverse clustering strategies enhanced binning performance, enabling the robust reconstruction of both cellular genomes and plasmids.

### Evaluating PlasMAAG plasmid binning using hospital sewage samples long-read data, and short-read plasmidomics data

Validating PlasMAAG binning performance on real data is not straightforward as current tools do not provide quality estimates for plasmids and might show inherent biases when exploring understudied environments such as wastewater. We instead applied a binning validation strategy based on sequencing both short– and long-read metagenomics from the same set of samples (**Fig. 4.A**). We considered a long-read contig to be composed of a set of short-read contigs if they aligned with 97% identity and a long-read contig coverage of 90% (see Methods). By tallying the number of such sets of short-read contigs binned together, we got a measure of recall of short-read contig binning. We observed that PlasMAAG community-based bins reconstructed 21% more long-read contigs than VAMB, the second-best performing binner (**Fig. 4.B**). Superior PlasMAAG binning performance was consistent even when accounting for incompleteness of the long-read assembled contigs (**Supplementary Note 5**, **Supplementary Figure 18**). To identify the subset of long-read contigs that originated from plasmids, we sequenced samples after a plasmid enrichment to obtain paired metagenomics and ‘plasmidomics’ samples as done previously (41) (see Methods). Long-read contigs were defined as plasmid contigs if they were either (1) at least 50% covered by plasmidomics reads or (2) circular and below 500 kb. We identified short-read contigs as originating from plasmid if they aligned well to any long-read contig identified as plasmid (see Methods). Using this criteria, PlasMAAG community-based reconstructed 138 NC plasmids, which was 33% more plasmid long-read fragments than the second best binner VAMB, and 431% more NC plasmids than SCAPP cycles (**Figure 4.B**). These results were consistent with the performance validated using unfiltered long-read contigs, demonstrating PlasMAAG’s robust binning capacity across diverse biological entities.

**Figure 4.**
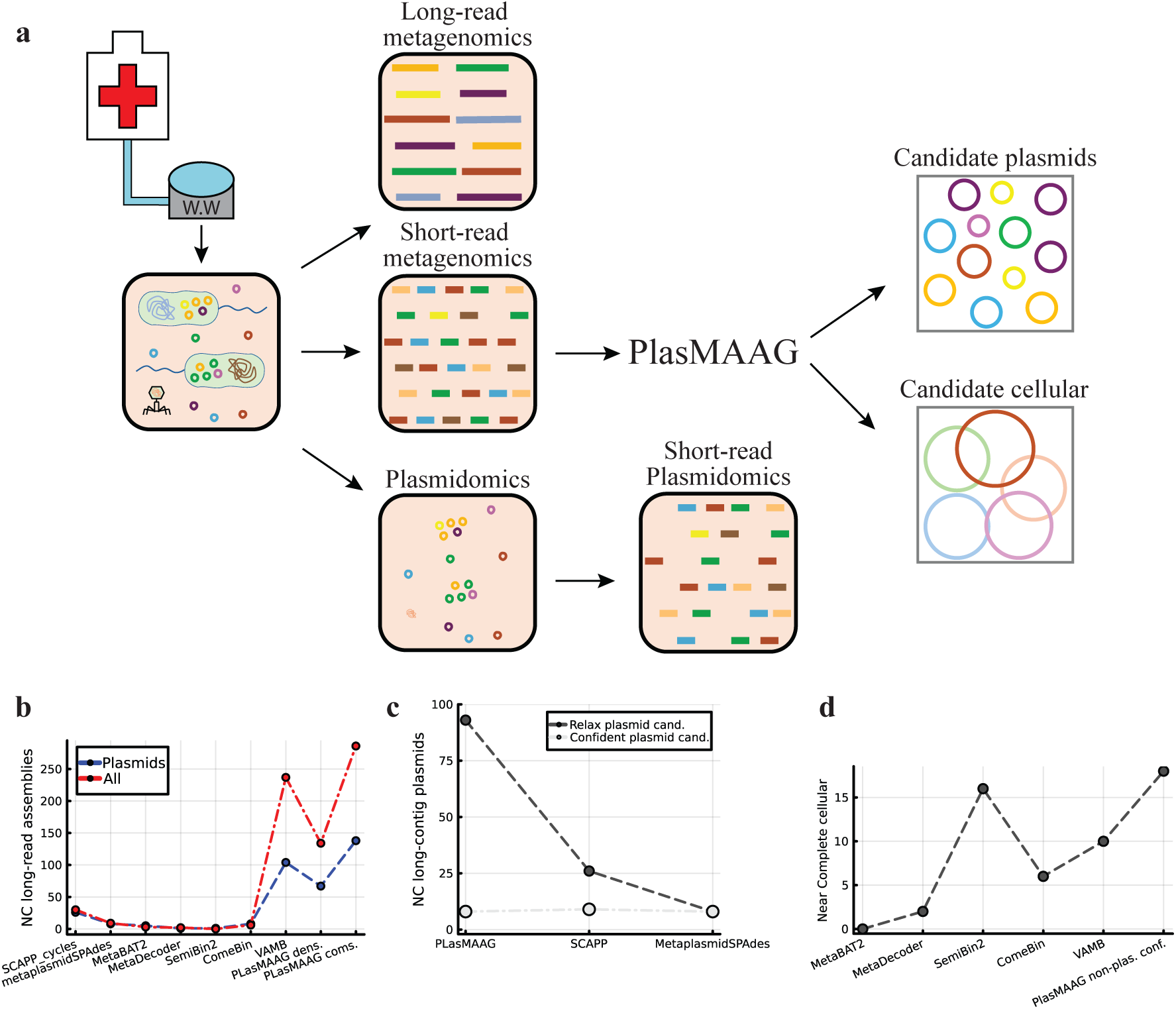
PlasMAAG on real samples from hospital sewage. **a**. Overview of the strategy used to validate PlasMAAG on the five hospital sewage samples. For each sample, long-read metagenomics, short-read metagenomics, and short-read plasmidomics datasets were generated (see Methods). PlasMAAG was applied to the short-read metagenomics data to produce candidate plasmid and cellular bins. These bins were validated against a reference assembly composed by long-read contigs to assess overall binning performance, and against a second reference assembly constructed from long-read contigs with plasmid evidence, identified either by circularity or plasmidomics read coverage. **b**. Binning performance of all methods across the five sewage samples, evaluated using all long-read contigs (red) and long-read contigs with plasmid evidence (blue). PlasMAAG dens.: bins produced using VAMB’s density-based clustering algorithm on PlasMAAG’s latents. PlasMAAG coms.: bins generated using the community-based clustering algorithm. **c**. Binning performance of PlasMAAG, SCAPP, and MetaPlasmidSPAdes under relaxed (light gray) and strict (dark gray) plasmid filtering criteria. **d**. NC cellular bins according to CheckM2 estimates, produced by all organism binners for the five hospital sewage samples. PlasMAAG non-plas. conf.: PlasMAAG density-based bins after extracting candidate plasmid contigs by aggregating geNomad plasmid contig scores per PlasMAAG community-based clusters (see Methods).

### Identifying plasmids in PlasMAAG bins using aggregated geNomad scores

When applying PlasMAAG to a real dataset with thousands of bins and no ground truth, we need to define a threshold to determine whether a bin contains a plasmid. This threshold balances precision and recall. To aid in this decision, we aggregated geNomad’s contig plasmid scores across all contigs within each bin. With a low threshold of 0.1, PlasMAAG reconstructed 93 NC long-read contigs highly confident plasmid based on the metaplasmidomics reads (long-read plasmidomics, LR-P), which represented a loss of 33% compared to not filtering with aggregated geNomad scores (**Figure 4.C**). Using a stricter geNomad plasmid threshold of 0.95 reduced the number of reconstructed LR-P to 8, a decrease of 94% (**Figure 4.D**). This implied that most long-read contigs, where we had experimental plasmid evidence, were predicted by geNomad to be of virus or chromosomal origin, as they had assigned a relatively low plasmid score (**Figure 4.B, Figure 4.C, Supplementary Figure 19**). By comparing aggregated geNomad scores with experimental plasmid evidence, we found that this mismatch mainly occurred where plasmid evidence was strong but not definitive (**Supplementary Note 6**, **Supplementary Figure 19-21**). This contrasted with the consistency observed in synthetic benchmarks, where geNomad generally demonstrated strong plasmid predictive performance (**Supplementary Figure 16**). Finally, we investigated the effect of this on cellular genomes and when applying a geNomad threshold of 0.95, the PlasMAAG density-based bins, which are the ones not classified as plasmid, were evaluated with CheckM2. We found 18 NC organisms, 3 more than SemiBin2, the second best binner on this dataset, and 8 more than VAMB. We found noticeable that PlasMAAG offered a better performance compared to SemiBin2, even though SemiBin2 leveraged single-copy genes whereas PlasMAAG did not. We conclude that PlasMAAG’s has state of the art performance on real datasets, both for reconstructing plasmids and cellular genomes.

### PlasMAAG enabled host-plasmids exploration from hospital sewage environments

By reconstructing plasmid and cellular genomes from the same samples, PlasMAAG enables an integrated analysis. We investigated host-plasmid abundance correlations of 24 hospital sewage samples collected in Spain (see Methods). PlasMAAG produced 27,954 candidate plasmid bins, and 213,431 non-plasmid candidate bins. PlasMAAG plasmid bins were aggregated into 13,912 cross-sample clusters, and bacterial hosts per plasmid cluster were inferred from PLSDB (see Methods). We identified 323 High quality cellular organism bins (HQ, completeness ≥ 70%, contamination ≤ 10%) and aggregated these using PlasMAAG cross sample cluster information. We found several significant positive correlations between candidate plasmid and cellular organism bins, for example, cluster *cl_20*, annotated as belonging to the *Aeromonas* genus, correlated with up to 41 plasmid clusters (adjusted p-value < 0.05), 12 of which were previously reported as known host-plasmid associations in the PLSDB database (**Figure 5A**). On the other hand, cluster *cl_293*, annotated as *Ruminococcus_E* genus, correlated with 43 plasmid clusters, none of them previously reported in PLSDB (**Figure 5A**).

**Figure 5.**
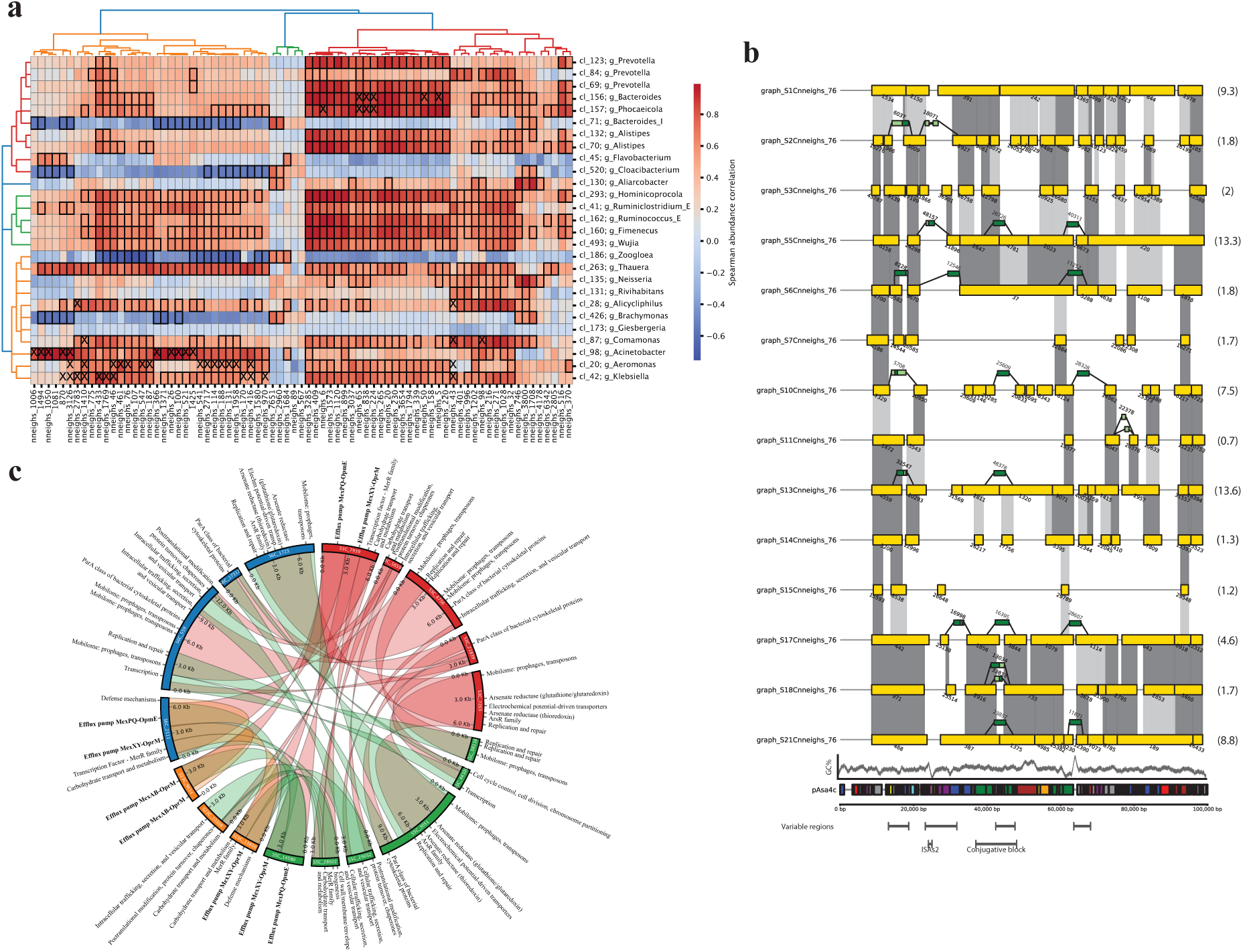
PlasMAAG enables host-plasmid association studies and exploration of intra-plasmid variation across environments, demonstrated using 24 hospital sewage samples. **a**. Spearman correlation between PlasMAAG high-quality (HQ) cellular clusters and PlasMAAG plasmid clusters with an aggregated geNomad plasmid score above 0.75. Highlighted cells with bold rectangles indicate significant correlations after Benjamini-Hochberg FDR correction. Cells marked with “X” represent plasmid-organism associations previously reported in PLSDB. The organism cluster dendrogram was generated using GTDB-tk taxonomic annotations, while the plasmid cluster dendrogram was based on abundance correlations. **b**. PlasMAAG plasmid cluster *nneighs_416* bins. Each row represents a bin from one sample, and numbers within parenthesis indicate median bin depth. Yellow blocks denote contigs aligned to *pAsa4c*, sorted by alignment position. Dark green blocks represent contigs not mapping to *pAsa4c* (see Methods), with their positions inferred from matches to other PLSDB plasmid accessions. Light green sections withing dark green blocks indicate alignment segments to *pAsa4c*. Dark grey areas indicate alignment graph edges, and light grey areas represent non-restrictive alignment matches (see Methods). GC%: Average GC content computed using a 1000 kb window. Colour code for pAsa4c regions: Blue (Replication and maintenance), Green (Conjugative transfer), Purple (Recombination and DNA repair), Orange (Secretion and surface structures), Red (Metabolism), Yellow (Enzymes), Cyan (Regulatory proteins and transcription factors), Brown (Transposases and mobile genetic elements), Gray (Hypothetical or unclassified). **c**. PlasMAAG plasmid cluster *nneighs_76*, composed of contigs from sample 6 (blue), sample 3 (red), sample 5 (green), and sample 23 (orange). Links represent alignment regions, coloured according to the sample of origin. Bold annotations indicate functions associated to antimicrobial resistance.

### PlasMAAG revealed intra-plasmid variation across hospital sewage samples

In PlasMAAG, contigs from different samples are projected into a shared latent space, enabling them to be clustered together, and split into per-sample bins thereafter. Aggregation of bins into PlasMAAG clusters enabled investigation of highly related plasmids from different samples. Cluster *nneighs_76* was selected for more in-depth analysis. Plasmid bins from the *nneighs_76* cluster reconstructed a 90 kb region from the plasmid *pAsa4c*, which is reported to be hosted by *Aeromonas salmonicida subsp*. (42) (**Figure 5B**). Despite representing highly overlapping regions of the same accession, bins from the *nneighs_76* cluster exhibited varying degrees of contig fragmentation. For instance, the bin from sample 2 was composed of 20 contigs, whereas the bin from sample 1 consisted of 10 contigs, which could be explained by the difference in the contig abundance. We then explored the relationship of the plasmid bins using the alignments from the AAG (**Figure 5B**). We also found that some bins in *nneighs_76* contains contigs that did not align to *pAsa4c.* Some of these unaligned contigs were found in multiple bins and were syntenic across bins aligned to each other, suggesting that we found true plasmid variation, and not an error in binning (**Figure 5B**). Using synteny, we could find four approximate locations on the reference sequence where these contigs belonged to. Three of four regions had hallmarks of recombination hotspots, including an ISAs2 insertion site, a known conjugative block and a segment with distinct GC content (42) (**Figure 5B**). Furthermore, 14 of 19 contigs not mapping to *pAsa4c* aligned to plasmid accessions reported to be hosted by organisms from the *Aeromonas* genus. Additionally, PlasMAAG clusters, together with the assembly-alignment graph, enable the exploration of diversity among similar plasmids across samples without PLSDB support. As an example, bins from the plasmid cluster *nneighs_416* exhibited a high degree of sequence similarity despite variations in contig fragmentation (**Figure 5C**). PlasMAAG facilitates the tracking of highly similar plasmids across different environments, allowing for the capture of their composition variations.

## DISCUSSION

Plasmids are pivotal in horizontal gene transfer, playing an influential role in shaping microbial communities. Their prevalence across microbial ecosystems highlights their importance, yet studying plasmids from environmental samples has been challenging due to their dynamic and unstable composition. This limitation has hindered efforts to bin and identify plasmids accurately, despite their abundance. The recent decrease in sequencing costs has significantly increased the availability of metagenomic samples, presenting an unprecedented opportunity to uncover plasmid diversity. However, the challenges of plasmid binning emphasize the need for a robust and broad-range plasmid binning method.

In this study, we introduced PlasMAAG, a novel deep learning-based framework for metagenomic binning of both plasmids and organisms. PlasMAAG leverages a unique feature we developed—assembly-alignment graphs—which enables the aggregation of assembly graphs across multiple samples. This advancement allows PlasMAAG overcome traditional limitations associated with single-sample plasmid assemblers.

PlasMAAG outperformed SCAPP, the current state-of-the-art plasmid assembler, on both synthetic and real datasets, delivering superior results for plasmid binning while being significantly faster. Besides producing more plasmid bins, the set of candidate plasmids produced by PlasMAAG achieved a more balanced trade-off between precision and recall, enabling a broader characterization of metagenomic samples. Notably, PlasMAAG’s capability to bin all sequences, including plasmids and organisms, offers a comprehensive approach to metagenomic analysis. PlasMAAG achieves organism binning results that are comparable to leading organism binners on synthetic datasets while demonstrating superior performance in understudied, real-world environments.

PlasMAAG’s holistic approach enables integrated studies, such as the exploration of plasmid-host associations. Using its comprehensive binning capabilities, we gathered correlation-abundance-based evidence for 773 plasmid-host associations, with only 7% previously reported in the PLSDB database. Furthermore, PlasMAAG’s assembly-alignment graph-based clustering revealed intra-plasmid variation across samples, enabling the study of plasmid sequence variation across environments.

We demonstrated that geNomad plasmid predictions were significantly enhanced when aggregated across PlasMAAG communities, underscoring the value of binning for refining plasmid sequence identification. However, we also saw that geNomad was inaccurate when applied to understudied environments, as validated by experimental paired metaplasmidomics, long, and short-read data. These discrepancies highlight the need for more robust plasmid sequence identifiers capable of handling complex or uncharted environments.

The success of PlasMAAG is largely attributable to the assembly-alignment graph, a feature that complements assembly graph signals across samples in a multi-sample framework. This innovation not only enhances binning accuracy but also facilitates the inference of compositional similarities between samples. Moreover, assembly-alignment graphs also improve binning of contigs from the same sample, through indirect links to contigs of other samples.

Another notable innovation in PlasMAAG is its use of contrastive loss to integrate traditional binning features like k-mer composition and contig abundances, with the assembly-alignment graph. This approach could be extended to incorporate other graph-like data in the binning process, such as Hi-C data. As sequencing technologies advance and contigs become decreasingly fragmented, particularly in long-read datasets, the utility of using cross-sample alignments to bridge gaps in the assembly graphs will grow, covering larger genome fractions and providing richer insights.

Despite the advances introduced by PlasMAAG, plasmid binning remains a significant challenge, as evidenced by the lack of groundbreaking plasmid binners in recent years. This underscores the necessity of innovative approaches, like PlasMAAG, that address the complexities of plasmid diversity and recombination. By enabling the study of plasmids alongside organisms from highly complex samples, PlasMAAG expands our ability to explore microbial communities comprehensively. Its focus on plasmids—an often-overlooked but critical component of microbial ecosystems—enhances our understanding of their role in horizontal gene transfer and microbial community dynamics.

In conclusion, PlasMAAG represents a step forward in plasmid and organism binning from metagenomic samples. By incorporating assembly-alignment graphs and contrastive learning, it addresses longstanding challenges in plasmid binning while providing a framework for studying plasmid-host associations and microbial community dynamics. PlasMAAG offers a valuable tool for advancing our understanding of microbial ecosystems, with implications for environmental microbiology, public health, and biotechnology. PlasMAAG

## MATERIAL AND METHODS

### Overview of PlasMAAG

The inputs to the PlasMAAG pipeline are a set of reads per sample. Reads are assembled per sample with *metaSPAdes* v3.15.5 (43) creating an assembly graph and contigs for each sample. The contigs across all samples are concatenated together to create the contig catalogue. Reads are mapped to the catalogue with *minimap2* v2.24 (44) and *samtools* v1.18 (45), creating per-sample BAM files. The alignment graph is generated by aligning the contigs across samples with NCBI *blast* 2.15.0 (46). The assembly– and alignment graphs are merged into the assembly-alignment graph (AAG). *Fastnode2vec* v0.05 (37), an optimized version of node2vec, is used to embed local AAG context of each contig into an embedding space, from which communities of contigs with similar embeddings are extracted. The k-mer composition and abundance features of contigs are embedding using a variational autoender (VAE), where an additional loss term is added which penalizes distance between contigs of the same community. Using the VAE embedding, communities are expanded, merged, and purified. The *geNomad* (38) tool is used to separate plasmid from non-plasmid contigs: Communities of plasmid contigs are extracted as separate bins, whereas the rest contigs are extracted in bins using a clustering algorithm.

### Benchmark datasets

We based our benchmark dataset on the existing CAMI2 short-read human microbiome toy dataset, but had to modify the dataset to allow benchmarking of plasmids: First, the original dataset did not provide assembly graphs, so we assembled the reads and mapped the resulting contigs back to the CAMI2 source genomes to determine their origin, using *minimap2* and accepting hits with an identity > 97% and a query coverage > 90%. Because this approach initially led to many unmapped or ambiguously mapping contigs, we re-simulated the reads using *wgsim* (47) with zero sequencing errors, then assembled each sample using *metaSPAdes* without the use of error correction. Second, CAMI2 considered plasmids to be part of their cellular host genome with the same abundance, which would inhibit our abundance-based binning approach. We changed so that plasmids were separate genomes with an abundance proportional to host abundance times a Gaussian random variable, as done in (18). Finally, CAMI2 did not contain reads simulated from across the edges of the underlying circular sequences, which prevents assembly graph cycles and hobbles graph peeling-based approaches like that used by SCAPP. We made sure to include such reads.

### Assembly graph edge weighting

Assembly graphs were extracted from the *assembly_graph_after_simplification.gfa* file generated from metaSPAdes and converted into a NetworkX v3.4.2 (48) directed graph, with contigs represented as nodes, and links between segments in contigs represented as edges. To enrich the assembly graph signal for binning, graph edges were weighted with the *normalized linkage* metric, which is dependent on the number of links established between any segments from each pair of contigs, normalized by the length of the contigs. For a pair of contigs *c^i^, c^j^*, the number of links connecting those contigs *n_links_i j_*, and the contig lengths *l^c^,* normalized linkage is:

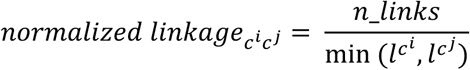

### Alignment graph edge weighting

After assembly, contigs shorter than 2000 bp were discarded as done in (26). Contigs were aligned all against all using NCBI blast using *blastn* command with *-perc_identity 95*, only keeping between-sample hits, alignment identity ≥ 98.0% and an alignment ≥ 500 bp. We also removed alignments between sequences that contained large sections that did not align due to sequence diversity, as we wanted the alignments to represent shared sequences across samples. The remaining set of alignments after filtering was defined as ‘restrictive’ alignments. From the aligments we created an alignment graph with contigs as nodes and alignments as edges. Edges were weighted with the *normalized alignment* metric to reflect the alignment certainty. For a pair of contigs *c^i^, c^j^*, alignment identity *id*, alignment length *L*, and contig length *l^c^:*

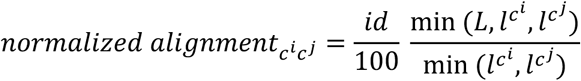

### Assembly-alignment graph community extraction with node2vec

Assembly and alignment graphs share no edges, since their edges connect only within-sample and between-sample contigs, respectively. This allowed us to trivially merge the graphs by adding the edges from one graph into the other, thus creating the AAG. To extract communities from the AAG, we first ran *fastnode2vec* on the AAG to obtain contig embeddings. We created a new graph by linking contigs within a cosine distance of 0.1 in embedding space, after which we defined each connected component to be a contig community. We optimized the *fastnode2vec* hyperparameters and clustering radius to generate pure communities at genome level, running a small grid search over the re-simulated CAMI2 Airways dataset. The embedding dimensions, walk length, number of walks, window size, p, and q parameters from fastnode2vec were set to 32, 10, 50, 10, 0.1, and 2.0. The embedding clustering cosine distance radius was set to 0.1.

### Contrastive-VAMB for community merging and expansion

Contrastive-VAMB is a variation of the original VAMB model, with a modification on the loss function to account for the communities extracted from the fastnode2vec embeddings. Contrastive-VAMB is composed of an encoder, latent representation layer m, and a decoder. Each contig represented by the concatenation of the contig co-abundances along samples **A**_in_, the tetranucleotide frequencies **T**_in_, and the unnormalized contig abundances **C**_in_ and passed to the encoder. The encoder projects the contigs into a latent normal N(μ, I) distribution parametrized by the m layer, from which the decoder samples. The decoder is optimized to reconstruct **A**_in_, **T**_in_, and **C**_in_ from the instances sampled from N(μ, I), decrease the latent cosine distance between contigs with closely related node2vec graph embeddings, and decrease the deviance between the latent normal distribution N(μ, I) parametrized by the μ layer and the standard normal distribution used as prior N(0, I).

### Loss functions

The contrastive-VAMB loss can be decomposed in three terms: reconstruction loss, contrastive loss, and regularization loss. The reconstruction loss (L_rec_) penalizes the reconstruction error of **A**_in_, **T**_in_, and **C**_in_. In the same way than the original VAMB reconstruction loss, cross entropy (CE) and sum of squared errors (SSE) losses were set for the reconstruction of the **A**_in_ and **T**_in_, respectively, whereas SSE loss was set for the **C**_in_ loss. These three terms are weighted with hyperparameters w_A_, w_T_, and w_C_.

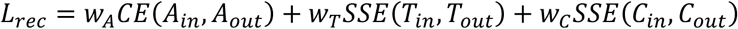

The contrastive loss (L_contr_) penalizes the cosine distance between the VAMB latent representations of the contigs and contigs highly related in node2vec embedding space, when such cosine distance overcomes a predefined margin *m*, *m* being a hyperparameter. For a contig *ci* and highly related fastnode2vec embedding space contigs *H^ci^* ={n_0,…,_ n_n_}:

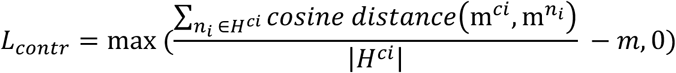

The regularization loss (L_reg_) penalizes the deviance between the latent normal distribution N(μ, I) parametrized by the μ layer, and the standard normal distribution used as prior N(0, I) with the Kullback-Leibler divergence, which since the standard deviation is set to 1, simplifies to:

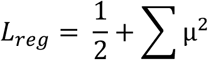

Finally, the model total loss (L) was aggregated with weighting hyperparameters *w_Lreg_*, and *w_Lcontr_*:

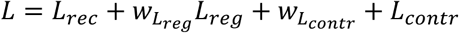

### Clustering plasmid/organism candidates with geNomad

Two parallel strategies were implemented to cluster the latent space tailored to extract plasmids and non-plasmids, respectively. The plasmid clustering strategy is composed of two phases: clustering community-based and clustering iterative medoid based, both based on latent space cosine distances. The clustering community-based works in five steps (**Supplementary Figure 9**): (1) for each community extracted from the node2vec embeddings, link contigs belonging to the same community, and remove links between contigs with a VAE embedding cosine distance > 0.2. (2) Contigs are recruited into the community if within 0.01 cosine distance to any community member. If the recruited contig is part of a community, the two communities are merged. (3) The expanded communities are extracted from the latent space as bins, and remaining contigs are clustered with the original medoid based VAMB clustering algorithm, (4) self-circularized contigs are extracted based upon mapping read-pairs where mates map to opposite contig ends within 50 bps from the contig end, and extracted from the clusters, (5) Plasmid score is defined for each cluster by aggregating the geNomad plasmid contig scores with a contig length weighted mean, defining plasmid candidates when cluster scores are larger than the defined threshold. When geNomad plasmid threshold is larger than 0.5, a fixed geNomad plasmid threshold of 0.5 is applied to the circular contigs, accounting for the circular evidence relatable to plasmids. The non-plasmid clustering strategy consists in 2 steps: (1) Cluster the VAMB-latent space with the iterative medoid clustering algorithm from VAMB. (2) Extract contigs belonging to any plasmid candidate cluster defined by the plasmid prone clustering strategy.

### Binning benchmarking – CAMI2 reassembled

We compared the plasmid and organism binning performance of PlasMAAG, VAMB v4.1.3, MetaBAT2 v2.12.1, SemiBin2 v2.1.0, Comebin v1.0.4, MetaDecoder v1.0.19, and SCAPP v0.1.4 over the re-simulated CAMI2 datasets. Binning performance was evaluated in terms of genomes recovered with precision ≥ 95% and recall ≥ 90%, so-called “NC genomes”. Since PlasMAAG, and VAMB, MetaBAT2, SemiBin2, Comebin, MetaDecoder perform the binning after assembling the contigs, precision and recall of the bins were obtained from the contig references, using BinBencher v0.3.0 (49). On the other hand, SCAPP and MetaPlasmidSPAdes v3.15.3 assemble their own contigs. Here, we produced a ground truth by aligning the output bins to the origin genomes using NCBI blast 2.15.0 accepting hits with an identity > 97% and a query coverage > 90%, after which we benchmarked using BinBencher.

### Sample benchmarking CAMI2 reassembled

Precision, recall, and F1 was computed for each set of plasmid candidates, reflecting the plasmid characterisation at the sample level, not at the bin level. Given a sample (*s*), a set of plasmid candidates (*candidates*), binning precision and binning recall thresholds (*pre*, *rec*), and the set of true plasmids present in the sample (*plasmids*):

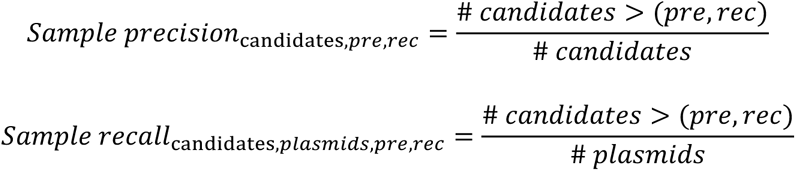

Enabling to compare the number of bins classified as plasmid, compared to the total number of plasmid genomes at specific binning precision and recall thresholds.

### Hospital sewage samples sequence datasets

Two datasets were used in this study to assess the quality of plasmid binning. Urban sewage samples (UWS) samples were collected from comparable UWSs from Denmark and Spain located in Odense and Santiago de Compostela, as previously described (41). In this study, only hospital sewage samples from each location were used. Sewage samples were collected in the winter and summer of 2018 using ISCO automatic samplers for 24-hour flow (50 mL per 5 min) in Denmark, while 24-hour-time proportional samples in SP (mixing hourly samples according to flow information) (**Supplementary Table 8**). Three replicates per site and season were collected on three consecutive days without rain events. All samples were initially cooled with ice on-site, then 100 mL of each sample was centrifugated at 10,000 g for 8 min at 4 °C in the laboratory. After removing supernatant, pellets were resuspended in 20 % of glycerol stock to reach a final volume of 10 mL for storage at −80 °C. In total, environmental DNA was extracted from all samples using NucleoSpin Soil kit (Macherey & Nagel, Dürein, DE) using 500μl of glycerol stock material for direct shotgun metagenomic using Illumina NovaSeq using 2×150bp paired-end mode (all samples) and PacBio Sequel2e (5 samples from Denmark). PacBio libraries were built from the same DNA extracts using libraries using SMRTbell express template 2.0 kit and Sequel II Binding Kit 3.2 (Pacific Bioscience, CA, USA) and barcoded using SMRTbell Barcoded Adapter Plate 3.0 (Pacific Bioscience, CA, USA). Two libraries per 8M SMRTcell (Pacific Bioscience, CA, USA) were pooled and sequenced on a PacBio Sequel2e instrument at University of Copenhagen.

For plasmids enriched samples, we used specific methods to deplete non-plasmid DNA as described previously (50, 51). Briefly, hospital sewage samples were pretreated by filtration, vortex and sonication and resuspended in TE buffer. Afterwards, a pre-lysis cocktail of cell-wall degrading enzymes: lysozyme, mutanolysin, and lysostaphin was used to facilitate lysis of Gram-positive bacteria during alkaline lysis. Pre-lysis was followed by alkaline lysis to remove chromosomal DNA (52), followed by Plasmid-Safe™ ATP-Dependent DNase (Lucigen, UK) digestion. Plasmid-Safe DNase will digest any fragments of dsDNA with open 3’ or 5’ termini, hence removing fragmented chromosomal DNA. The purified plasmid DNA was then quality-checked, libraries prepared and sequenced on an Illumina NextSeq platform with a v2.5 sequencing kit (Illumina, San Diego, CA, USA) in paired-end mode.

### Binning benchmarking – hospital sewage

We compared the binning performance of PlasMAAG, VAMB, MetaBAT2, SemiBin2, Comebin, MetaDecoder, metaplasmidSPAdes, and SCAPP over the 5 hospital sewage samples. Performance evaluation was based on the long-read sequences generated from the same samples and defined by the long-read contigs recovered with precision ≥ 95% and recall ≥ 90%, so-called “NC long-read assemblies”. To evaluate the overall binning performance, the entire set of long-read contigs was used to build the reference. Whereas to evaluate the plasmid binning performance, only the long-read contigs either circular or with metaplasmidomics reads coverage > 50% were used to build the reference. To build the references, we mapped the short-read contigs to either set of long-read contigs to determine their origin, using minimap2 v2.24 and accepting hits with an identity > 97% and a query coverage > 90%, and used Binbencher for the benchmarking. To account for plasmid circularity, 2 copies of each long-read contig were concatenated before mapping the short-read contigs. adovNC organisms were estimated with CheckM2 v0.1.3.

### Host-plasmid and intra-plasmid diversity exploration

PlasMAAG was used to bin the contig sequences from 24 hospital sewage samples from hospitals in Spain. PlasMAAG bins were aggregated into PlasMAAG clusters and classified as plasmids if the aggregated geNomad plasmid score exceeded 0.75, defining them as plasmid clusters. Only plasmids clusters with more than 150 kb were considered for the host-plasmid association. Organism’s bin quality was estimated with CheckM2 v0.1.3, and only high-quality (completeness ≥ 70% and precision ≥ 90%) (HQ) bins were kept. GTDBtk v2.4.0 (53) was used to estimate taxonomy for the HQ bins, with cluster taxonomy assigned based on majority vote. Abundance correlation analysis was only conducted for plasmids and organism’s clusters with non-zero abundance over at least 18 overlapping samples. Spearman correlation coefficients and p-values were computed using *scipy.stats.spearmanr*. To account for multiple testing, p-values were corrected using the Benjamini-Hochberg (FDR) correction implemented in the *statsmodels.stats.multitest.multipletests* package. Plasmid cluster hosts were inferred from PLSDB when aligning to any PLSDB entry with >80% identity and >80% coverage. Functional annotations of contigs were performed with *anvi’o* v8 software, using the ‘anvi-run-workflow-w contigs’ command.

### Resource usage

We evaluated computational resource usage of all methods using the Airways CAMI2 re-assembled dataset and five samples from the hospital sewage dataset. For the Airways dataset, PlasMAAG used 46 minutes, 8 threads, and 16 GB of RAM. In contrast, SCAPP, excluding the BAM file generation step, took 192 minutes, utilized 16 threads, and required 24 GB of RAM (**Supplementary Table 5**). Among the other binners, PlasMAAG was slower than VAMB, MetaDecoder, and MetaBAT2. For example, VAMB completed the task in just 8 minutes while using 8 threads and 16 GB of RAM. However, we observed a different trend when evaluating performances on the five hospital sewage samples. When accounting for the additional steps of read assembly and read mapping required to compute abundances, PlasMAAG exhibited similar runtimes to most binners, except for SCAPP, which required significantly more time. Specifically, PlasMAAG took 3,575 minutes, VAMB took 3,435 minutes, ComeBin required 4,911 minutes, and metaplasmidSPAdes took 4,430 minutes (**Supplementary Table 6**). In contrast, SCAPP required 116,965 minutes—32 times longer than PlasMAAG. This difference in runtime remained consistent even when excluding the read assembly steps (**Supplementary Table 6**).

## DATA AVAILABILITY

Reads, contigs, and contig annotations for the re-assembled CAMI2 datasets are available here: https://erda.ku.dk/archives/826fe4d8889f88db2ec20058f9eaa015/published-archive.html and https://erda.ku.dk/archives/fb2c6dd2a8e002becb58233bd4388f7c/published-archive.html. The metagenomic short reads, metaplasmidomic short reads, and metagenomic long reads from the 5 Danish hospital sewage samples, as well as the metagenomic short reads from the 24 Spanish hospital sewage samples, are available in the European Nucleotide Archive under BioProject PRJEB85938, whereas the assemblies for all samples are available here: https://erda.ku.dk/archives/e87f0d5e12ca4c1204379d4932c3ae59/published-archive.html (**Supplementary Table 8**).

## SUPPLEMENTARY DATA

Supplementary Data are available online.

## AUTHOR CONTRIBUTIONS

S.R. and S.J.S. conceived the study. S.R., J.N.N, and P.P.L guided the analysis. P.P.L. developed PlasMAAG, wrote the software, and performed the analysis. Additionally, J.N.N., and L.S.D. also wrote the software. J.N.N. also performed analyses. S.J.S, J.N, M.P.A, and L.J.J. provided input for the analysis. I.K and J.N generated the sewage data. P.P.L., J.N.N., and S.R. wrote the manuscript with contributions from all co-authors. All authors read and approved the final version of the manuscript.

## ACKNOWLEDGEMENTS

P.P.L., J.N.N., L.S.D, M.P.A., and S.R. were supported by the Novo Nordisk Foundation grant NNF23SA0084103. Furthermore, P.P.L., J.N.N., J.N., I.K., S.J.S., and S.R. were supported by the Novo Nordisk Foundation grant NNF20OC0062223. L.J.J. and S.R. were supported by the Novo Nordisk Foundation grant NNF14CC0001. M.P.A was funded by the Novo Nordisk Foundation Bioscience Ph.D. Program grant No. NNF22SA0078231. L.J.J. was supported by the Novo Nordisk Foundation grant NNF14CC0001. PacBio sequencing Facility at University of Copenhagen’s Biology Department is supported by Novo Nordisk Foundation grant NNF20OC0061528 to S.J.S.

## CONFLICT OF INTEREST

S.R. is the founder and owner of the Danish company BioAI and has performed consulting for Sidera Bio ApS. The remaining authors declare no conflict of interest.

## CODE AVAILABILITY

PlasMAAG is freely available at https://github.com/RasmussenLab/vamb/tree/vamb_n2v_asy/workflow_PlasMAAG.

## Notes

https://erda.ku.dk/archives/826fe4d8889f88db2ec20058f9eaa015/published-archive.html

https://erda.ku.dk/archives/fb2c6dd2a8e002becb58233bd4388f7c/published-archive.html

https://erda.ku.dk/archives/e87f0d5e12ca4c1204379d4932c3ae59/published-archive.html

### Summary of Updates

Figure 4.A updated from PLAMB to PlasMAAG.

