## Supplementary material for "Accurate plasmid reconstruction from metagenomics data using assembly-alignment graphs and contrastive learning"

### Supplementary Note 1: PlasMAAG plasmids balance precision-recall across the benchmark datasets

While reconstructing many plasmids is important, the overall quality of the candidate plasmid sets proposed by any method is crucial. This is particularly relevant because, unlike chromosomal sequences, estimating plasmid completeness and contamination with external tools is very challenging. To address this, we evaluated the candidate plasmid sets produced by SCAPP, geNomad, and the plasmids generated by averaging scores across PlasMAAG community-based clusters. These three methods have nevertheless fundamental differences that complicate direct comparisons. SCAPP relies on assembly graph cycles, so it is expected to perform well on fully assembled and well-defined plasmids within the assembly graph. However, SCAPP's approach primarily captures a subset of the plasmids present in the sample, providing precise characterization for only that plasmid subset. In contrast, geNomad is a pure contig classifier, which makes it more prone to generating incomplete plasmids while providing a broader, though more fragmented, characterization. Lastly, PlasMAAG, as a non-cycle-based binner, has the potential to reconstruct both highly assembled plasmids and partially assembled ones, offering a more comprehensive sample characterization that includes both complete and incomplete plasmids, acting as a balance between SCAPP and geNomad. To validate this hypothesis, and considering the properties of each method, we evaluated their ability to reconstruct medium-quality plasmids (precision >0.9, recall >0.5) per sample. Overall, PlasMAAG highly confident plasmids produced candidate plasmid bins with a better balance between precision and recall, achieving F1 scores that were 14-46% higher than SCAPP and 43-248% higher than geNomad along all datasets **(Figure 2.H).** In consistency with our hypothesis, SCAPP produced sets of plasmids that characterized the samples with higher precision but low recall for 4/5 datasets, whereas geNomad provided plasmid sets with higher recall for 4/5 datasets (**Figure 2.E**, **Supplementary Figure 1**). However, when evaluating PlasMAAG's performance using geNomad thresholds, the balance between precision and recall can be adjusted based on user priorities (**Supplementary Figure 1**). Those results corroborate that PlasMAAG achieves an optimal balance between precision and recall, providing a broad but precise plasmid characterization.

**Supplementary Note 2: GeNomad plasmid predictions were highly improved by binning insights**

Identifying sequences originated from plasmids is very challenging due to their capacity to acquire genetic material from taxonomically unrelated sources (1). On the other hand, it has been previously shown how binning can help to the contig classification since it groups contigs that potentially originated from the same source (2, 3). Considering the plasmid binning capacity of PlasMAAG, it offers great potential to contribute to the contig classification problem. GeNomad individual contig classification based on the aggregated sequence-based and markers-based branches yielded moderate PRAUCs and MCCs of 0.16-0.57 and 0.13-0.45 across the five benchmark datasets (**Figure 2.I-J**). When averaging the geNomad aggregated contig scores per PlasMAAG community-based clusters, an increase over the PRAUC (MCC) of 18-72% (28-106%) was achieved over all datasets, consistent with our hypothesis (**Figure 2.I-J**). We found that the score aggregation per cluster improvement to be grounded by the low level of organism-plasmid contamination in the PlasMAAG community-based clusters, where only 0.8-0.13% of bins in our benchmark datasets had more than 5% of the total bin size be both organism and plasmid (**Supplementary Table 1**). We then evaluated the geNomad sample composition estimation, which serves as the foundation for calibrating the geNomad scores. Our analysis revealed that geNomad tends to overestimate the plasmid composition in samples (**Supplementary Figure 2**). Given the inaccuracies in the sample composition estimation, and the circular nature of reusing scores as both input and output in the calibration process, we decided not to apply score calibration. Therefore, we believe that updating contig scores with binning information proved to be greatly beneficial for the plasmid sequence identification, improving the characterization of the samples.

**Supplementary Note 3: Plasmid/cellular genomes encoding differences on VAMB-contrastive latent space**

We found that plasmid and cellular genomes to be distributed differently in the latent space from VAMB-contrastive. The distinct latent space properties of plasmids and organisms were reflected by the difference in binning performance based on the clustering strategy. Plasmid binning was favored when extracting bins based on the community aggregation strategy, reconstructing 32-56% more NC plasmid bins than the ones using the density-based clustering from VAMB over 4/5 benchmark datasets (**Supplementary Figure 12**). On the other hand, we observed higher organism reconstruction when the same latent space was clustered with VAMB's density-based clustering algorithm, producing 72-238% more NC organism bins than the community-based approach over the benchmark datasets (**Supplementary Figure 12**. It is well established that plasmids have a great capacity to acquire and release genetic material through recombination compared to cellular genome (6, 18). The higher recombination of plasmids compromises the homogeneity of TNF and contig co-abundances along the plasmid sequence, hindering binning solely based on those features. Consistently with this plasmid property, we found a 15-62% higher overlap between the TNF distance distributions of intra-plasmids and inter-plasmid contig pairs, in comparison with the overlapping between intra-organism and inter-organism TNF distances distributions (**Supplementary Figure 13, Supplementary Table 4**). Additionally, we also found a 19-117% higher Pearson correlation between contig abundances from different plasmids, compared to the distribution of Pearson correlations between contig abundances from different organisms (**Supplementary Figure 13, Supplementary Table 4**), emphasizing the contribution of graph features for plasmid binning, which have proved to be key on previous attempts to reconstruct plasmid sequences (19, 20). Following the same trend, a 171-217% higher overlap between intra-plasmid and inter-plasmid contig pairs was observed over the latent space with respect the organism case, supporting distinct clustering approaches for organisms and plasmids (**Supplementary Figure 13, Supplementary Table 4**). We hypothesize the poor performance of community clustering for organisms may be due to the incompleteness of communities, which impacts organisms more, as their genomes are typically fragmented into more contigs than plasmids, which together with the insufficient single contig recruitment into those communities (**Supplementary Table 3**), and the observed high precision and relatively low recall (**Supplementary Figure 14**), generates organism bins with highly incomplete organism genomes. To summarize, TNF and abundance features behave differently in plasmids and organisms. These differences impact the latent space arrangement of their respective contigs and justify the use of custom clustering strategies.

**Supplementary Note 4: geNomad plasmid threshold effect on plasmid and organism classification from PlasMAAG latent representations**

Given the differences between plasmid and organism latent representations in PlasMAAG (**Supplementary Note 3**), we applied a clustering strategy to account for these variations leveraging geNomad plasmid predictions. This approach was sensitive to the geNomad threshold used for the classification. For instance, when the geNomad threshold was set to 0.1, 66-100% of the NC organisms bins were misclassified as plasmid. On the other hand, plasmids were less sensitive, since for the strictest plasmid threshold of 0.9, only 18-42% NC plasmids were misclassified (**Supplementary Figure 15**). The higher threshold sensitivity observed for organism bins can be explained by the less precise prediction of organisms when aggregating geNomad scores per cluster, compared to plasmids (**Supplementary Figure 16**).

**Supplementary Note 5: PlasMAAG leads binning reconstruction from waste-water hospital samples even when accounting for reference fragmentation**

Considering that long-read contigs only represent fragments of the original genomes present, the superior performance may be attributed to the fragmentation of the original genomes, which could advantage high-precision binners over high-recall binners. In other words, if binners reconstruct the entire genome and not only the long-read contig fragments, using a fragmented references would assign low precision to those bins, though it would still assign high recalls. Thus, we evaluated binners performances at a fixed recall threshold of 0.9, and decreasing precision thresholds of 0.95, 0.8, 0.6, 0.3, and only for bins composed by more than one contig, in that way compensating the binning evaluation for the potential fragmentation of the references. We did not observe substantial changes in the binning performance for all remaining binners along the decrease of the precision thresholds, whereas PlasMAAG community-based showed an increase of 531% bins along the decreasing precisions. These findings suggested that none of the binners except PlasMAAG community-based were binning contigs from the same fragments, nor from the same original genomic entities (**Supplementary Figure 18**).

**Supplementary note 6: GeNomad plasmid scoring conflicts with meta-plasmidomics evaluation**

Considering the decrease on the number of PlasMAAG NC long-read plasmid contigs when filtering bins with geNomad (**Supplementary Figure 19**), we investigated the congruity of geNomad scoring and the metaplasmidomics long-read filtering strategy. Namely, we investigated the distribution of geNomad plasmid scores for long-read contigs with decreasing plasmid evidence: (1) long-read circular contigs at least 95% covered by metaplasmidomics reads, and shorter than 500kb, (2) long-read circular contigs shorter than 500kb, (3) long-read linear contigs covered by metaplasmidomics reads at least by 95%, and shorter than 500kb, (4) long-read linear contigs at maximum 10% covered by metaplasmidomics reads, and longer than 100kb (**Supplementary Figure 20**). We found both strategies to agree on the extreme cases, where geNomad assigned high plasmid scores to long-read circular contigs at least 95% covered by metaplasmidomics reads, and shorter than 500kb as plasmids, and very low plasmid scores to long-read linear contigs at maximum 10% covered by metaplasmidomics reads, and longer than 100kb. However, conflicting evidence was found for the intermediate cases, which represents 89% of all long-read contigs. We propose two possible explanations causing this mismatch: (I) leakage of virus reads during the metaplasmidomics reads generation, (II) geNomad underscoring plasmid sequences, due to the high complexity environment. However, for (1) and (2) sets of long-read contigs, the distribution of geNomad virus scores does not support that the long-read contigs belonging to those sets to mainly originated from virus, since for both cases, 75% of the contigs had a virus score lower than 0.25 (**Supplementary Figure 21**).

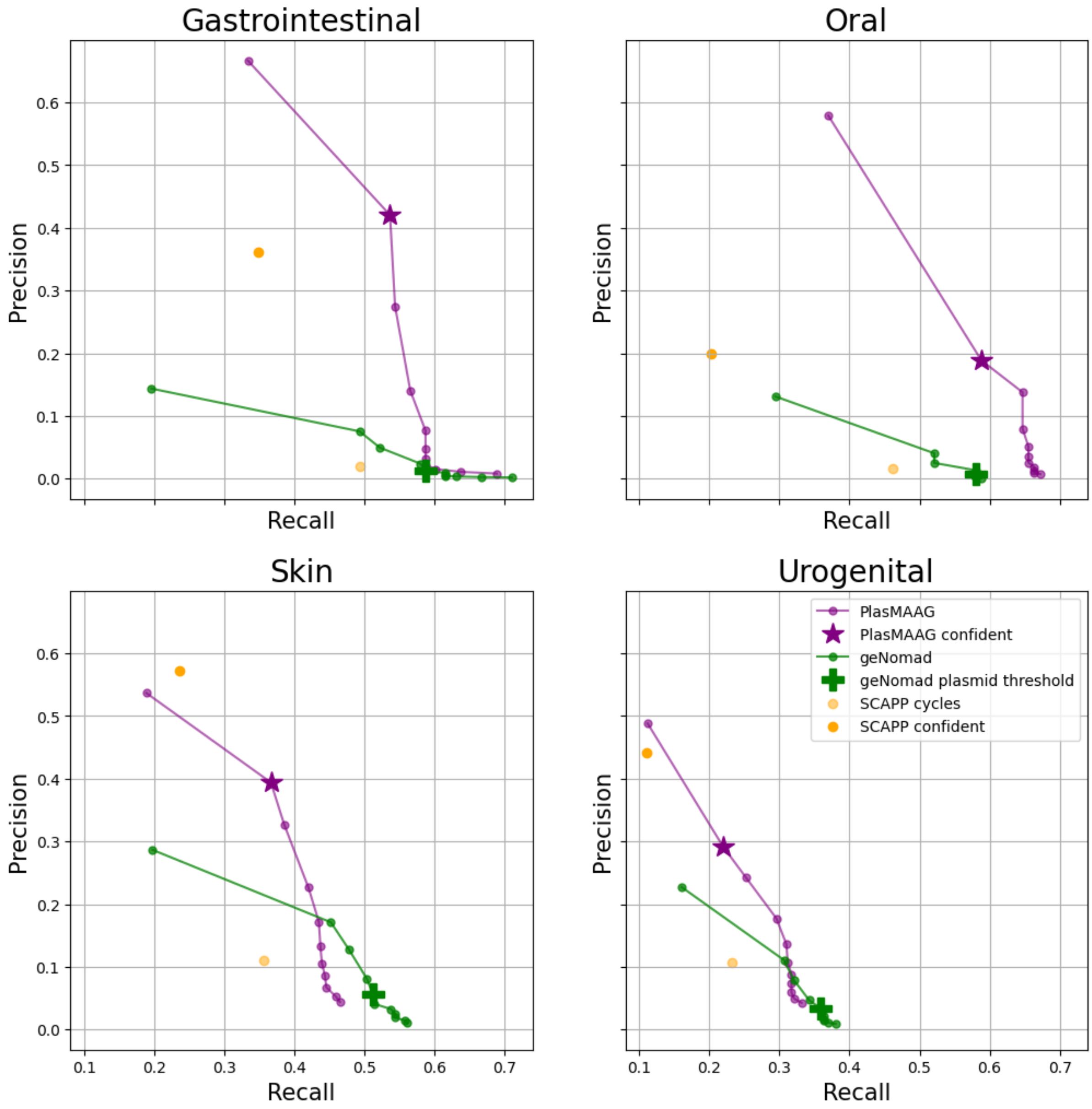


**Supplementary Figure 1. Sample recall and precision of plasmid candidates reconstructed by SCAPP, geNomad, and PlasMAAG across all CAMI2 re-assembled datasets except Airways.** Sample recall represents the fraction of medium-quality (MQ) plasmids reconstructed relative to the total plasmids present per sample, based on each method's proposed candidates (see Methods). Sample precision represents the fraction of actual plasmids among all candidate plasmids proposed by each method at the MQ level (see Methods). For geNomad and PlasMAAG, precision and recall were calculated at increasing geNomad thresholds of 0.1, 0.2, 0.3, 0.4, 0.5, 0.6, 0.7, 0.8, 0.9, and 0.95.

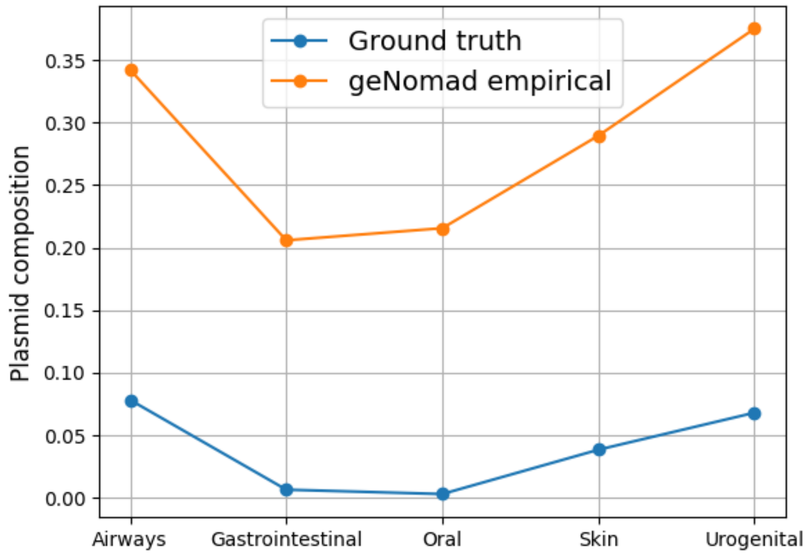


**Supplementary Figure 2. Empirical plasmid sample composition estimated by geNomad against real plasmid composition along the CAMI2 re-assembled benchmark datasets.** For each benchmark dataset, the *ground truth* composition was computed as the fraction of contigs mapping to plasmids, whereas *geNomad empirical* is the composition estimated by geNomad based on the geNomad scoring.

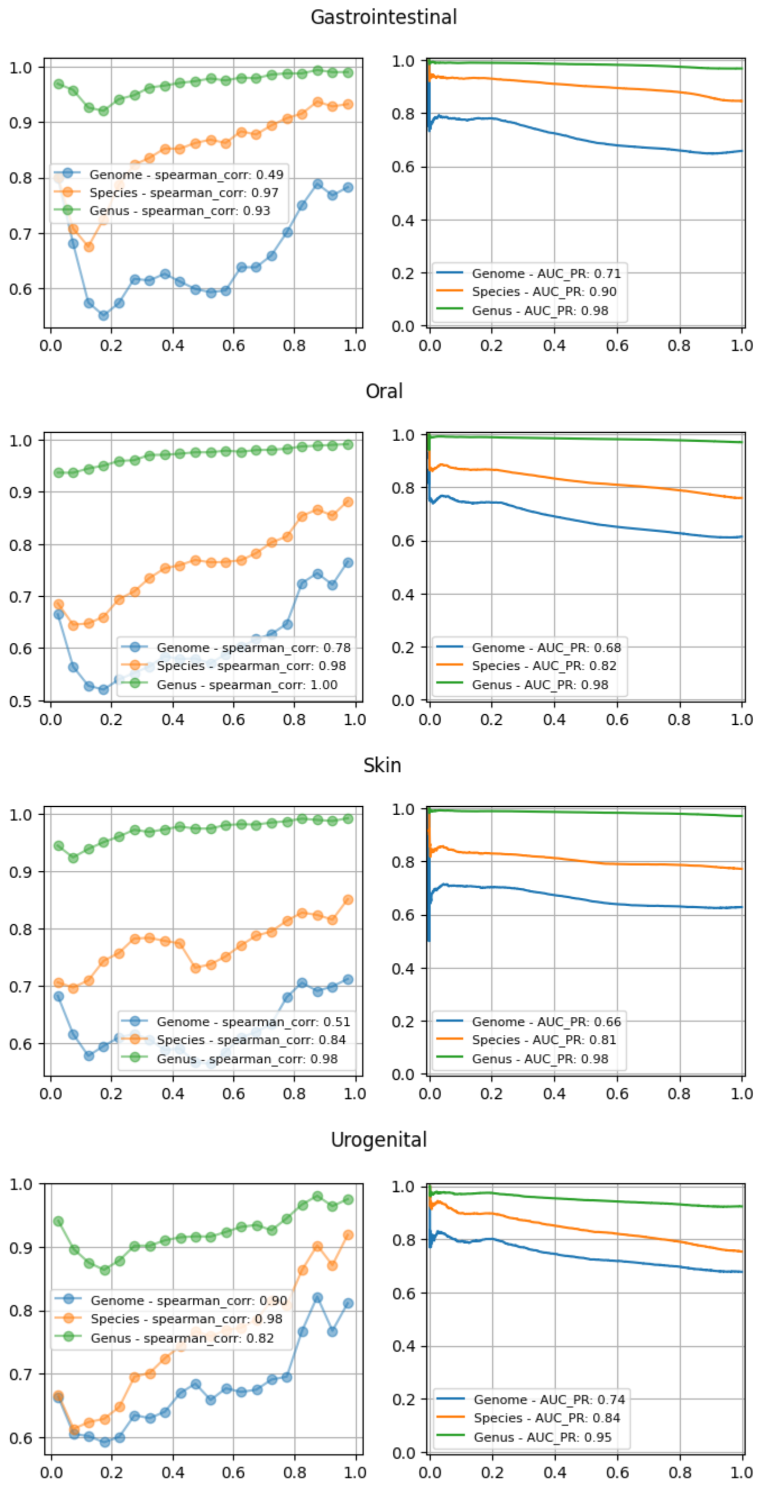


**Supplementary Figure 3**. **Average precision of assembly graph edges, sorted by weight and grouped into 5% bins (left), and precision-recall curve of the assembly graph edge weights.** Left column: Average normalized linkage against average precision, at genome (blue), species (orange), and genus (green) taxonomic levels. Right column: Precision recall curve of the normalized linkage weighting scheme, at genome (blue), species (orange), and genus (green) taxonomic levels. 140,556 69,767

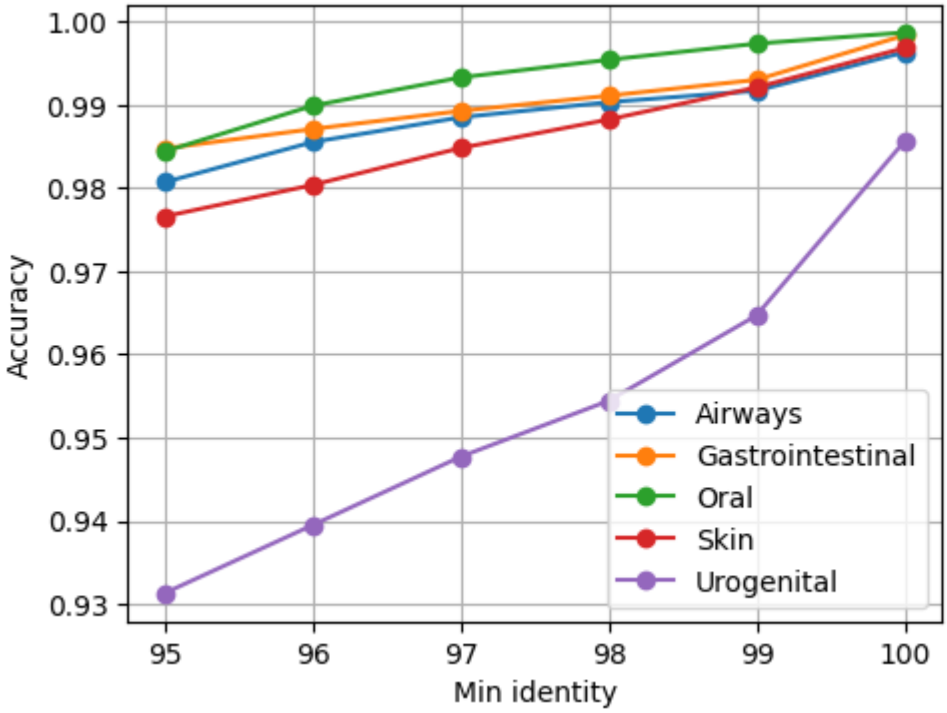


**Supplementary Figure 4**. **Alignment accuracy at increasing identity thresholds.** Alignment accuracy at the species level across increasing identity thresholds, with minimum alignment coverage set to 500 bps, and when only restrictive alignment is allowed (see Methods).

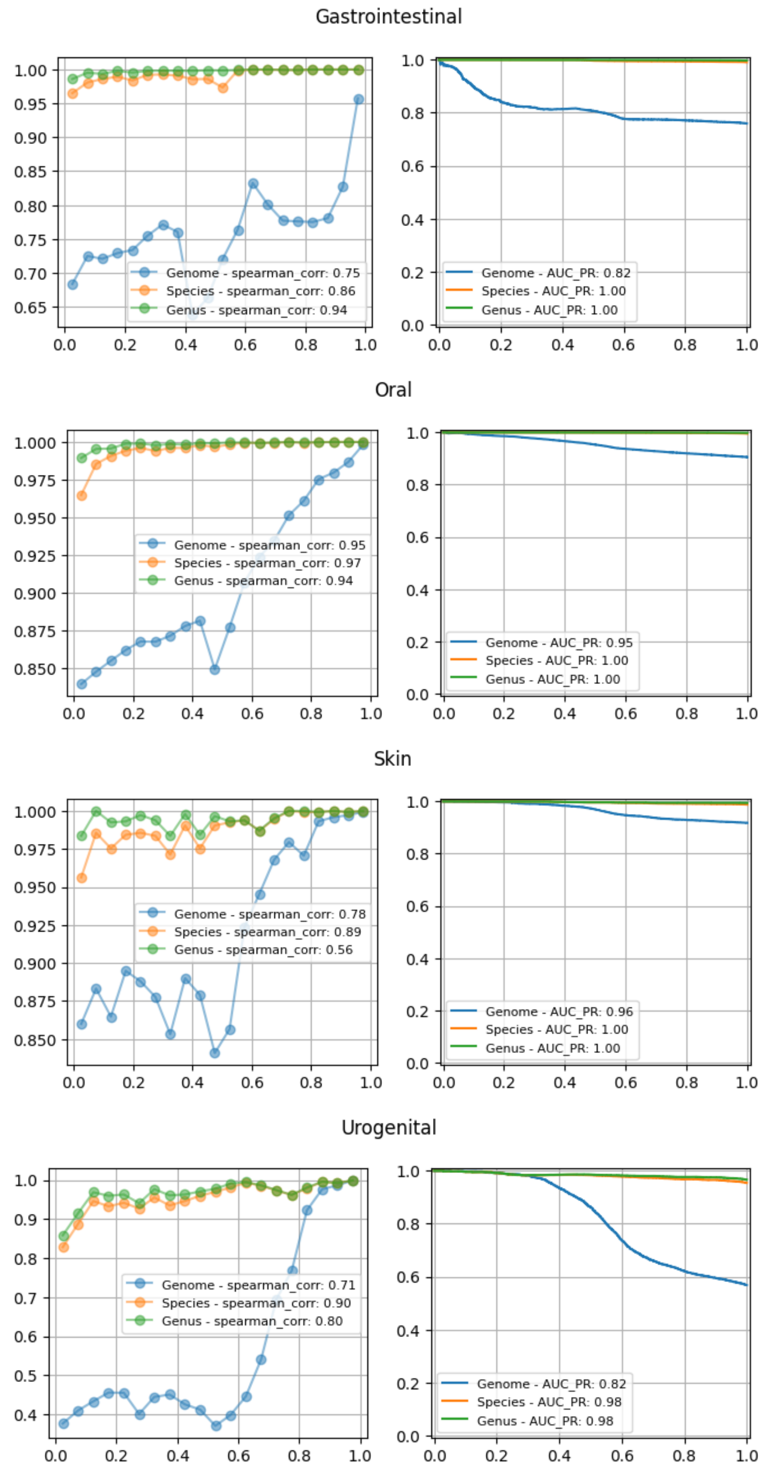


**Supplementary Figure 5**. **Alignment graph normalized alignment metrics across the CAMI2 re-assembled benchmark datasets.** Left column: Average alignment graph edge weights against average precision, at genome (blue), species (orange), and genus (green) taxonomic levels. Right column: Precision recall curve of the normalized alignment weighting scheme from the alignment graph at genome (blue), species (orange), and genus (green) taxonomic levels.

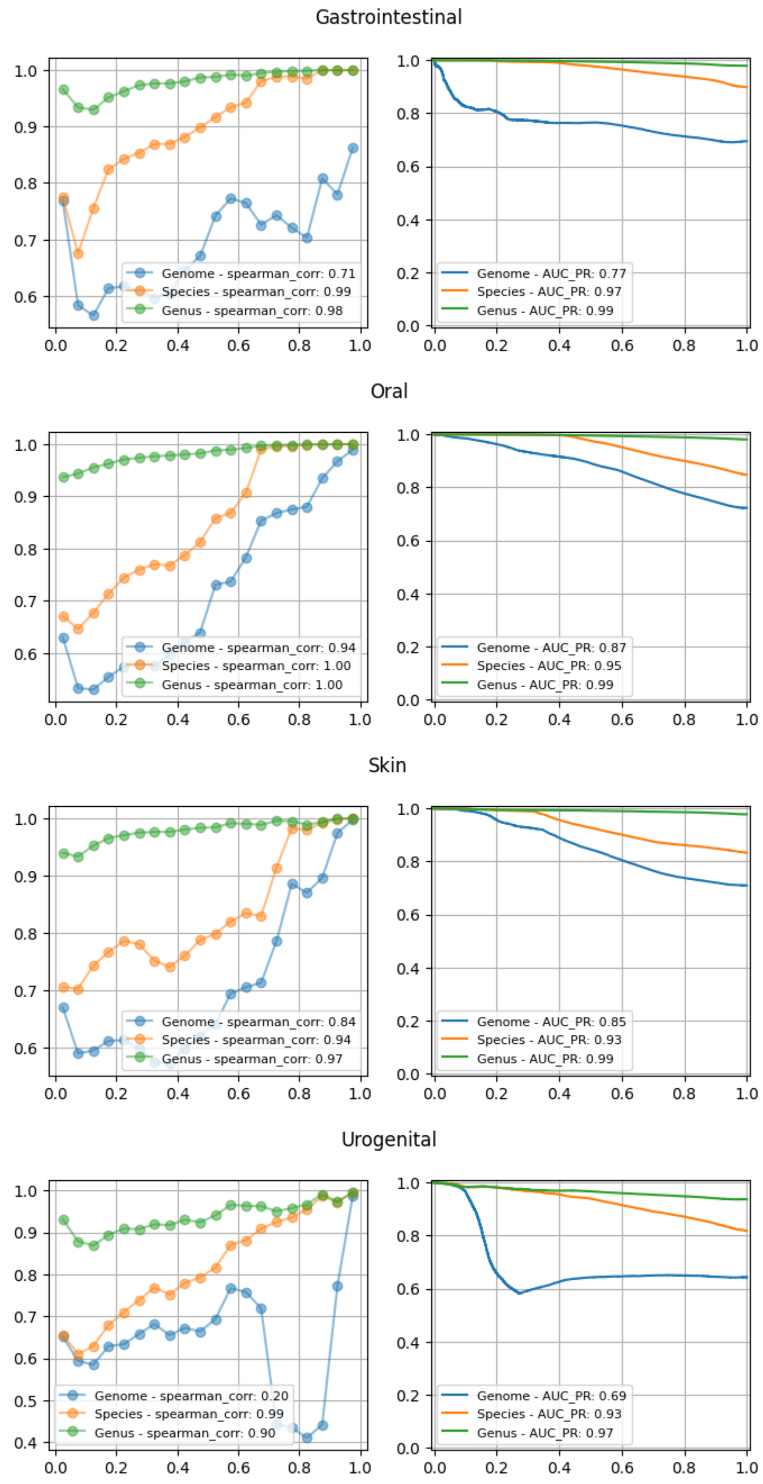


**Supplementary Figure 6. Assembly-alignment graph edge weighting metrics across the CAMI2 re-assembled benchmark datasets.** Left column: Average assembly-alignment graph edge weighting against average precision, at genome (blue), species (orange), and genus (green) taxonomic levels. Right column: Precision recall curve of the edge weighting scheme at genome (blue), species (orange), and genus (green) taxonomic levels.

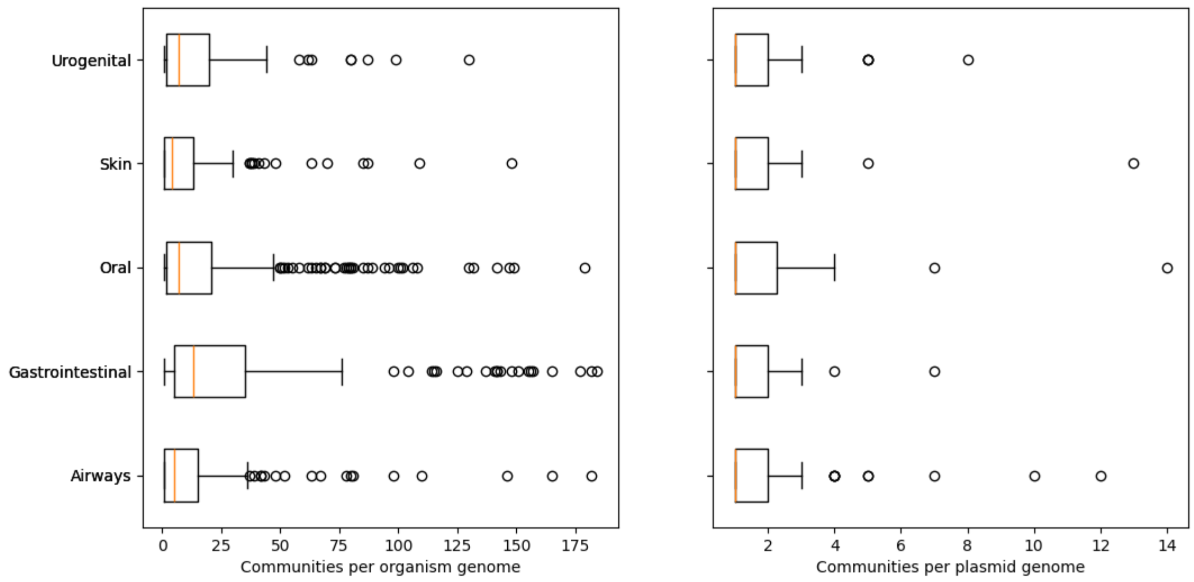


**Supplementary Figure 7. Organism and plasmid genome fragmentation according to the communities extracted from fastnode2vec assembly-alignment graph embeddings.** Communities were extracted from fastnode2vec assembly-alignment graph embeddings and evaluated based on the number of communities each organism (left) and plasmid (right) were split into. Results are presented for the re-assembled CAMI2 benchmark datasets.

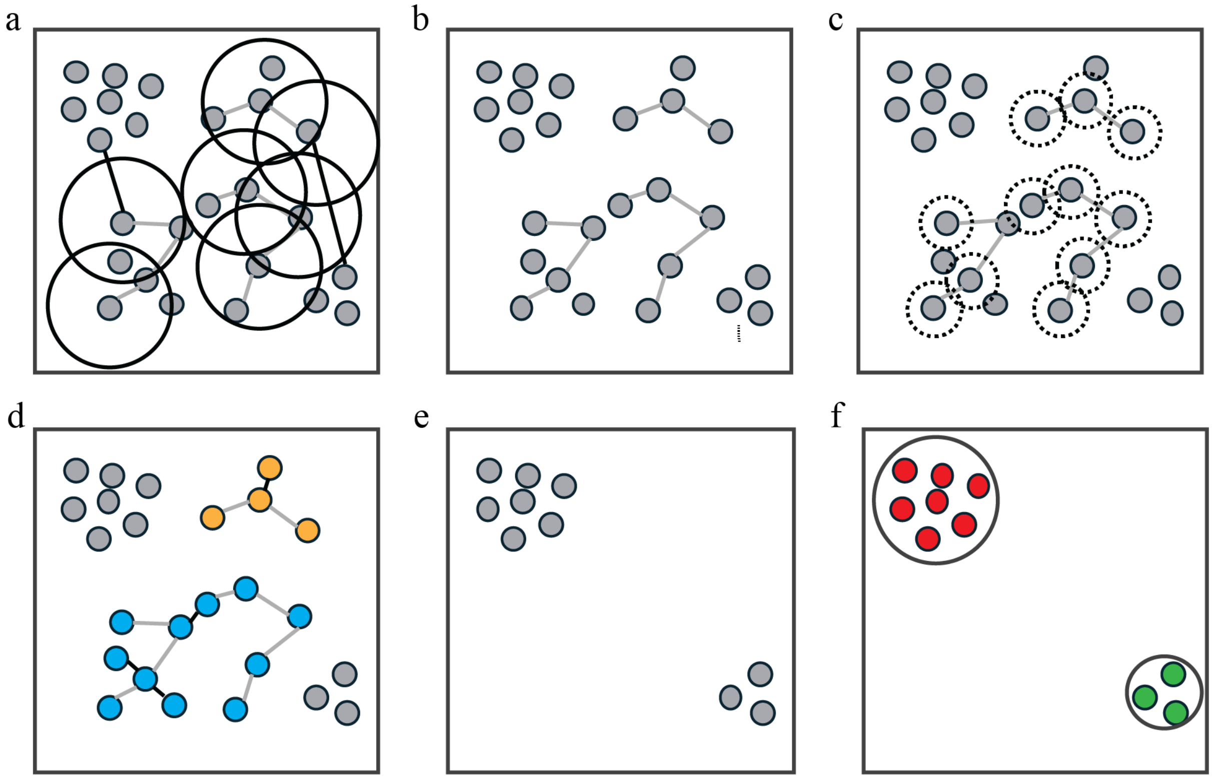


**Supplementary Figure 8. PlasMAAG community-based clustering algorithm. a**. Latent space distances between contigs from the same community are evaluated, identifying contig edges with distant latent representations (black edges), which overcome a predefined radius distance (black circles). **b**. Edges between contigs from the same community but with distant representations are removed. **c**. The latent proximity of contigs belonging to a community is scanned, identifying additional contigs that will be recruited into the communities if lie within the proximity radius (dashed circles). **d**. Contigs within the proximity latent radius are recruited into the community. **e**. Split and expanded communities are withdrawn from the latent space. **f**. Remaining contig latent representations are clustered with the default VAMB clustering algorithm.

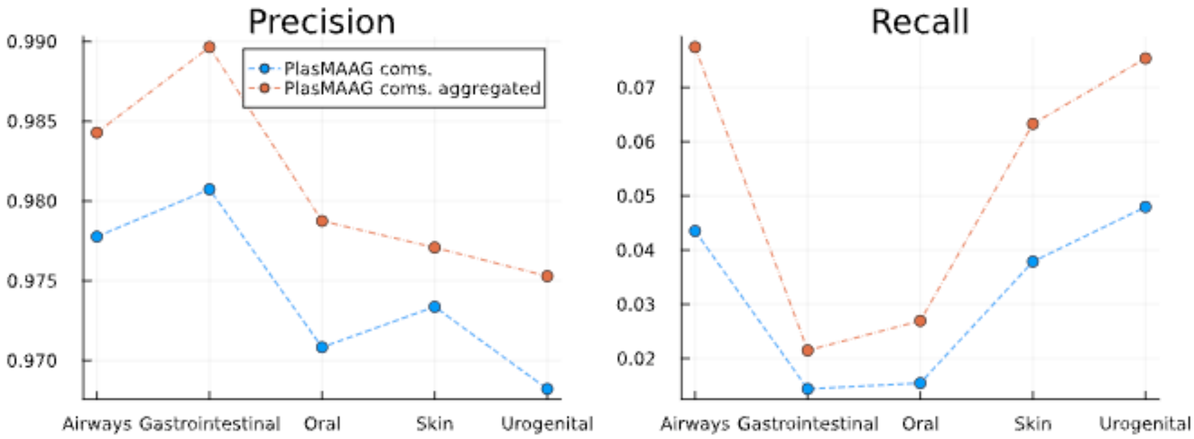


**Supplementary Figure 9. Impact of PlasMAAG's community-based clustering on the precision and recall of reconstructed bins.** Precision and recall values are shown before (blue) and after (red) merging, expanding, and splitting communities using VAMB-contrastive across the re-assembled CAMI2 benchmark datasets. PlasMAAG coms.: Communities extracted from the assembly-alignment graph embedding; PlasMAAG coms. aggregated: clusters extracted from PlasMAAG latent space when applying the community-based clustering (see Methods).

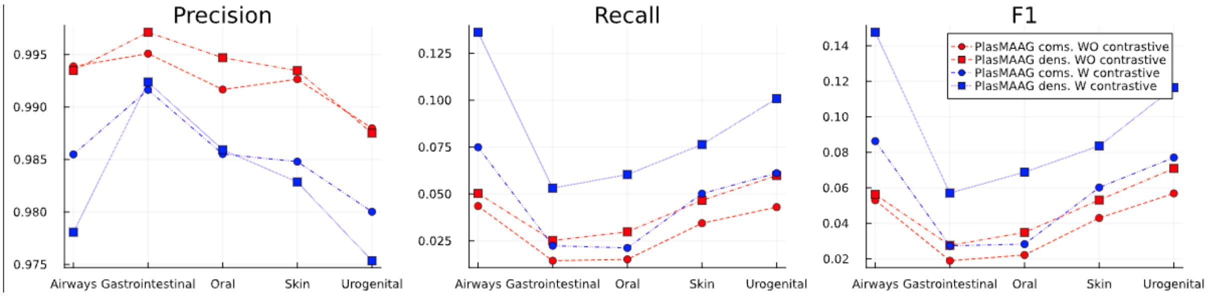


**Supplementary Figure 10. Impact of contrastive loss on PlasMAAG's community-based and density-based clusters, evaluated using precision, recall, and F1-score across the re-assembled CAMI2 benchmark datasets.** PlasMAAG community-based (circles) and density-based (squares) bins precision, recall, and F1 when running VAMB with (blue) and without (red) contrastive loss. PlasMAAG coms.: clusters extracted from PlasMAAG latent space when applying the community-based clustering; PlasMAAG dens.: clusters extracted from PlasMAAG latent space when applying the density-based clustering; W contrastive: when latent representations were generated with VAMB-contrastive; WO contrastive: when latent representations were generated with default VAMB.

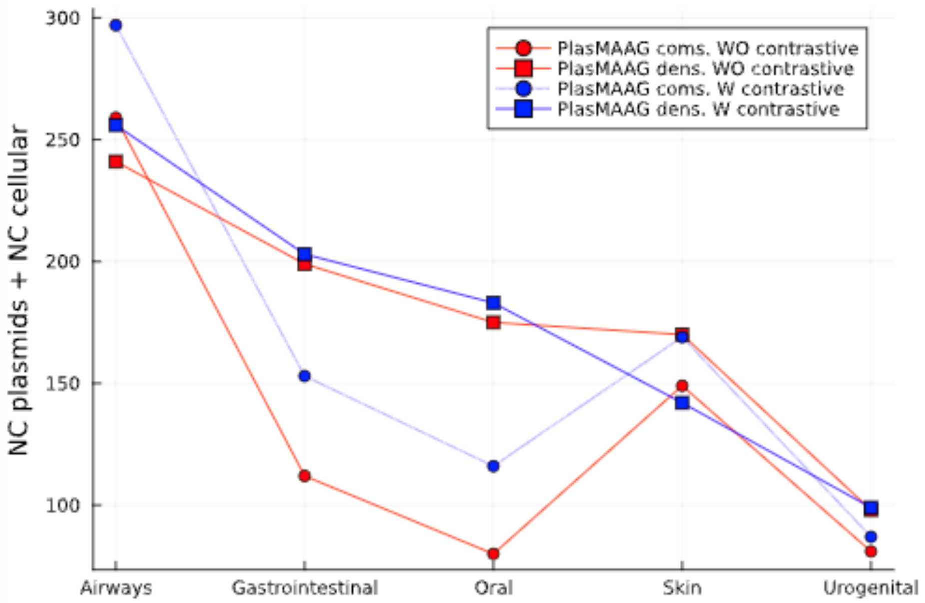


**Supplementary Figure 11. Impact of contrastive loss on PlasMAAG's community-based and density-based clusters, assessed by the number of NC bins reconstructed from the re-assembled CAMI2 benchmark datasets.** Precision, recall, and F1 values for PlasMAAG community-based (circles) and density-based (squares) bins are shown when running VAMB with (blue) and without (red) contrastive loss. airways: Airways; gi: Gastrointestinal; oral: Oral; skin: Skin; urog: Urogenital. PlasMAAG coms.: clusters extracted from PlasMAAG latent space when applying the community-based clustering; PlasMAAG dens.: clusters extracted from PlasMAAG latent space when applying the density-based clustering; W contrastive: when latent representations were generated with VAMB-contrastive; W contrastive: when latent representations were generated with default VAMB.

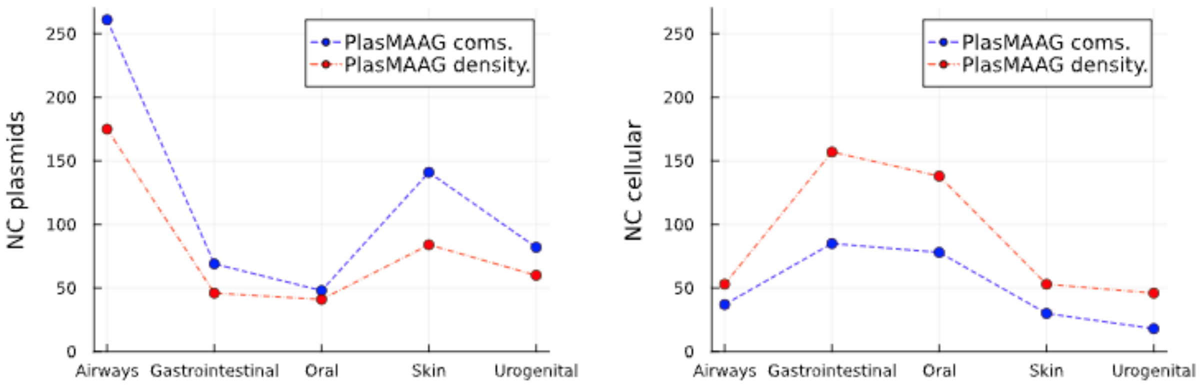


**Supplementary Figure 12. Plasmid and cellular reconstruction performance of PlasMAAG's community-based and density-based clustering strategies across the re-assembled CAMI2 benchmark datasets.** The figure shows the number of near-complete (NC) plasmids (left) and NC cellular (right) reconstructed by PlasMAAG community-based (PlasMAAG coms.) and PlasMAAG density-based (PlasMAAG density.) clustering.

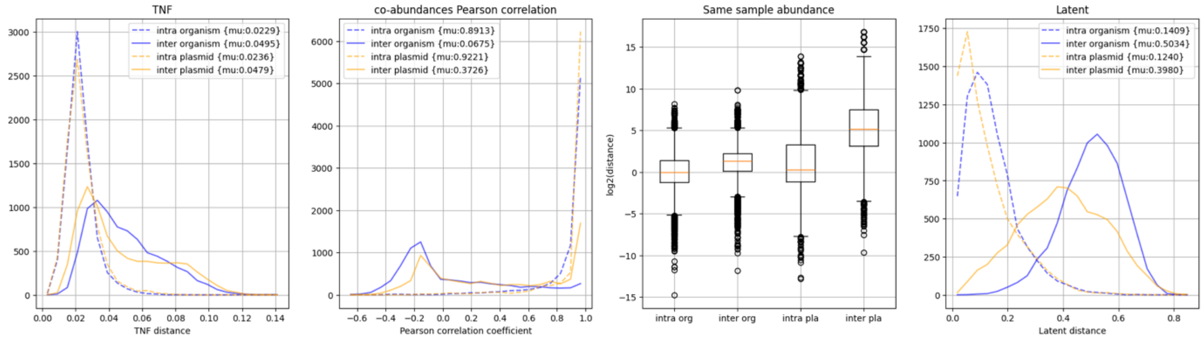


**Supplementary Figure 13. Plasmid and organisms have distinct signatures.** Distribution of TNF, co-abundance Pearson correlation coefficient, logarithm of the absolute intra-sample abundance, and latent distances, between pairs of contigs. For each curve or boxplot, 8000 contig pairs were sampled, which satisfied each condition: same cellular organism (intra organism), different cellular organism (inter organism), same plasmid (intra plasmid), inter plasmid (different plasmid). Sampled pairs of contigs were not repeated and were enforced to belong to the same sample. Contigs sampled from the re-assembled CAMI2 Airways dataset.

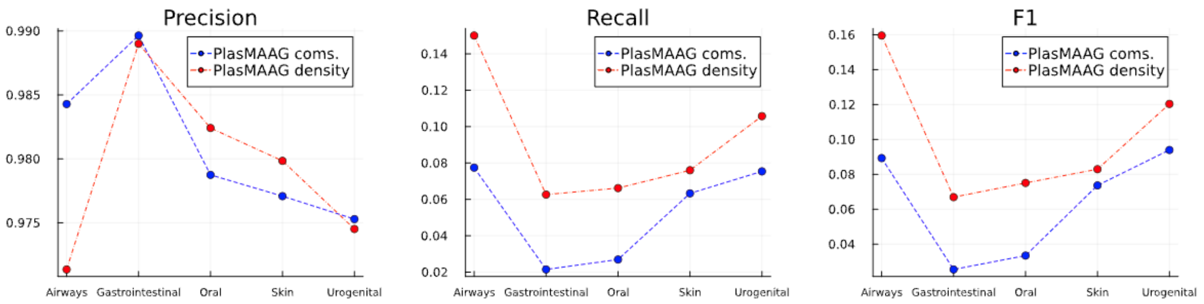


**Supplementary Figure 14. Precision, recall, and F1 of PlasMAAG community-based and density-based clusters across the re-assembled CAMI2 benchmark datasets.** Precision, recall, and F1 of bins generated with the community-based (PlasMAAG coms.) and density-based (PlasMAAG density) clustering algorithm.

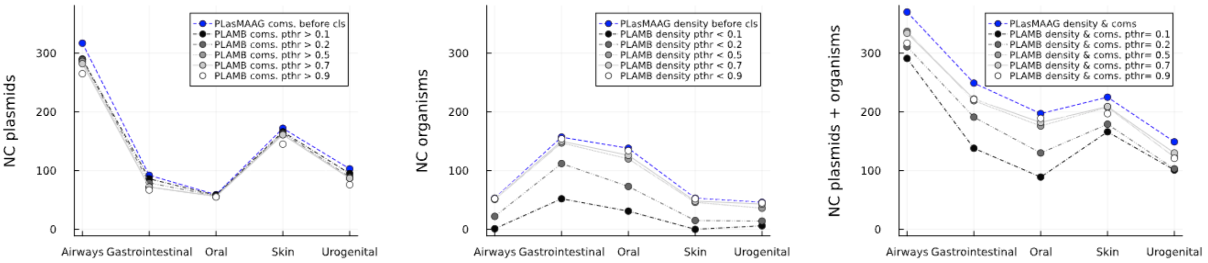


**Supplementary Figure 15. Near complete plasmid, organism, and plasmids and organisms when applying increasing geNomad filtering thresholds across the re-assembled CAMI2 benchmark datasets.** Near-complete plasmids (left), cellular (center) and plasmids + cellular (right) after classifying PlasMAAG community-based clusters as plasmids based on increasing geNomad plasmid thresholds. For each PlasMAAG community-based cluster that had an average plasmid bin score above the threshold, contigs belonging to such cluster were subtracted from the PlasMAAG density-based clusters (see Methods). Blue curves represent the absolute NCs before classification. pthr: aggregated geNomad plasmid threshold; PlasMAAG coms. before cls: PlasMAAG community-based clusters before using geNomad to select candidate plasmids; PlasMAAG density before cls: PlasMAAG density-based clusters before extracting plasmid clusters.

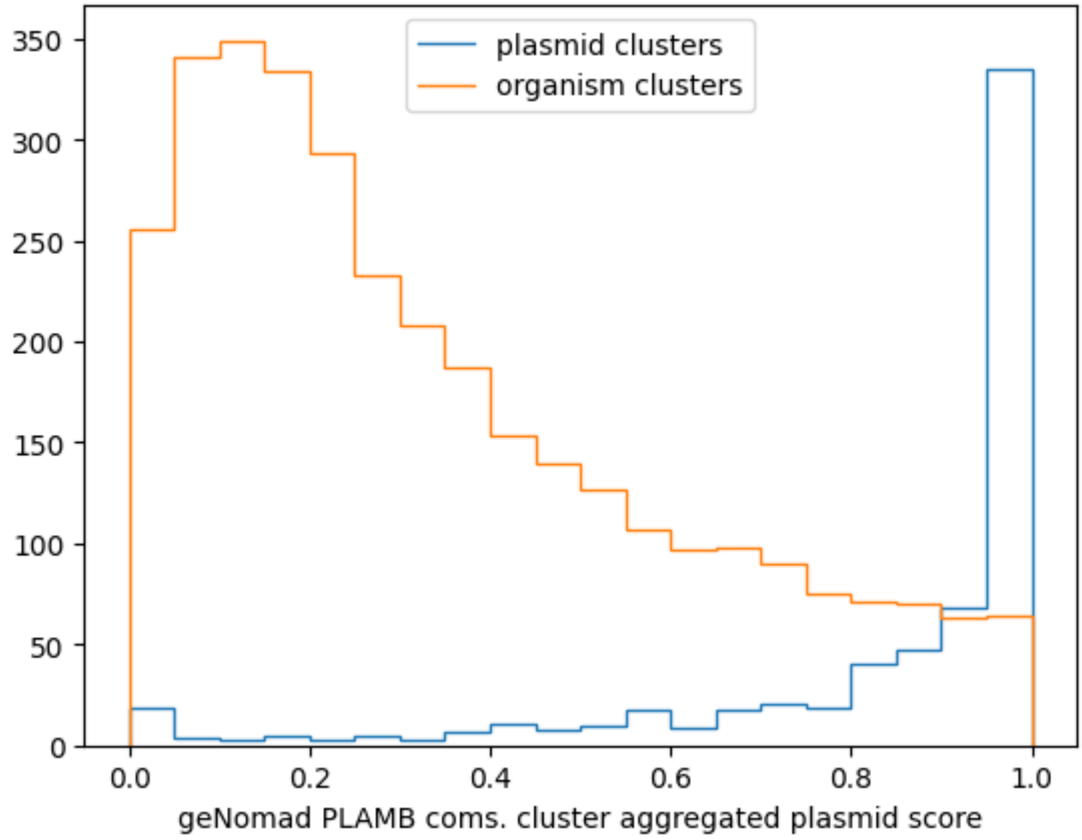


**Supplementary Figure 16. Plasmid and cellular organism geNomad score distributions of PlasMAAG community-based clusters.** Distribution of plasmid contig scores when averaging geNomad plasmid scores per PlasMAAG community-based clusters, for plasmid and cellular organism clusters. Results shown for the clusters generated from the Airways CAMI2 re-assembled dataset.

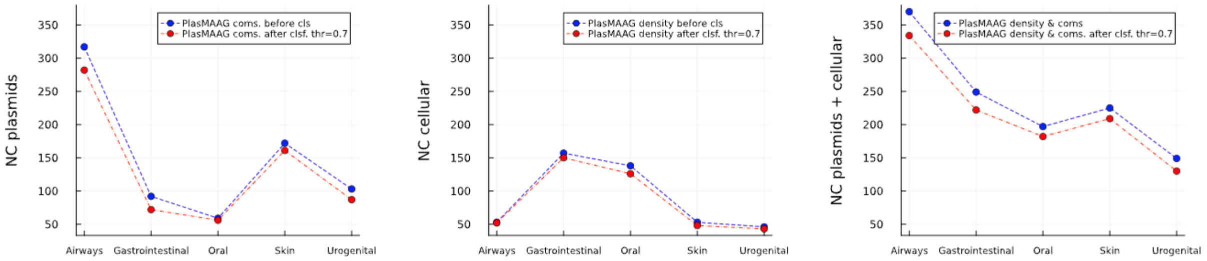


**Supplementary Figure 17. Remaining NC plasmids, cellular organisms, and plasmids + cellular organisms after filtering clusters based on geNomad scores.** Near-complete plasmids (left), cellular organisms (center) and plasmids + cellular organisms (right) before (blue) and after (red) classifying PlasMAAG community-based clusters as plasmids if average geNomad plasmid score greater than 0.7. For each PlasMAAG community-based cluster that had an average plasmid bin score above 0.7, contigs belonging to such cluster were subtracted from the PlasMAAG density-based clusters. airways: Airways; gi: Gastrointestinal; oral: Oral; skin: Skin; urog: Urogenital. PlasMAAG coms.: clusters extracted from PlasMAAG latent space when applying the community-based clustering; PlasMAAG dens.: clusters extracted from PlasMAAG latent space when applying the density-based clustering; before cls: clusters before using geNomad based classification into plasmid and cellular organism bins; after cls: clusters after using geNomad based classification into plasmid and cellular organism bins.

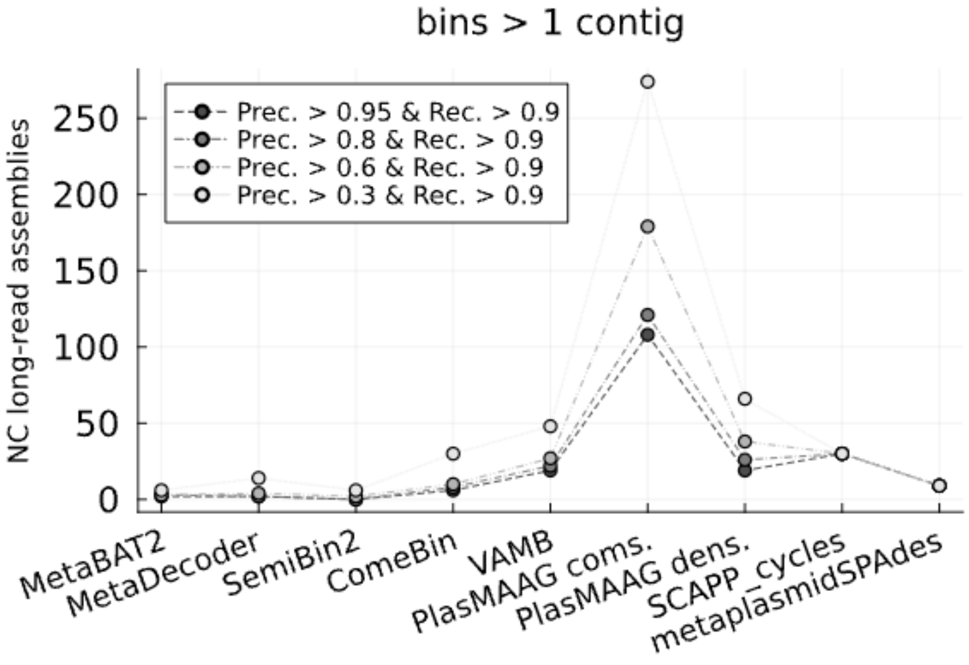


**Supplementary Figure 18. NC long-read contigs at decreasing precision thresholds reconstructed from the 5 hospital sewage samples.** Near-complete long-read contigs along decreasing precisions and fixed recall for all binners. Prec: Precision; Rec: Recall; bins > 1 contig: only bins containing more than one contig considered.

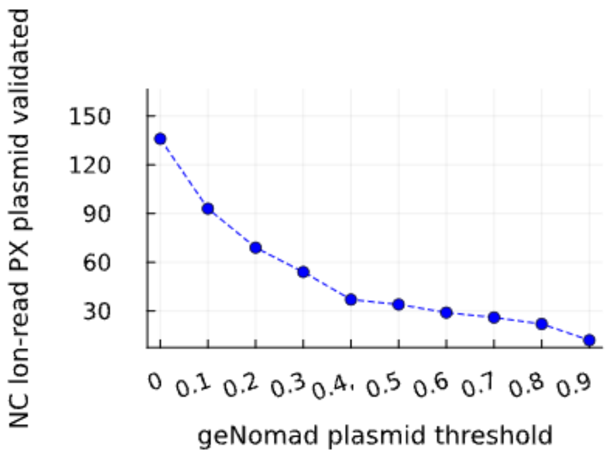


**Supplementary Figure 19. NC long-read plasmid contigs reconstructed by PlasMAAG from the 5 hospital sewage samples after applying geNomad filtering.** Near-complete long-read plasmid contigs reconstructed by PlasMAAG community-based bins, along increasing geNomad plasmid thresholds. Long-read contigs were considered plasmid if at least 50% covered by plasmidomic reads, or circular and below 500 kb. 79,6239

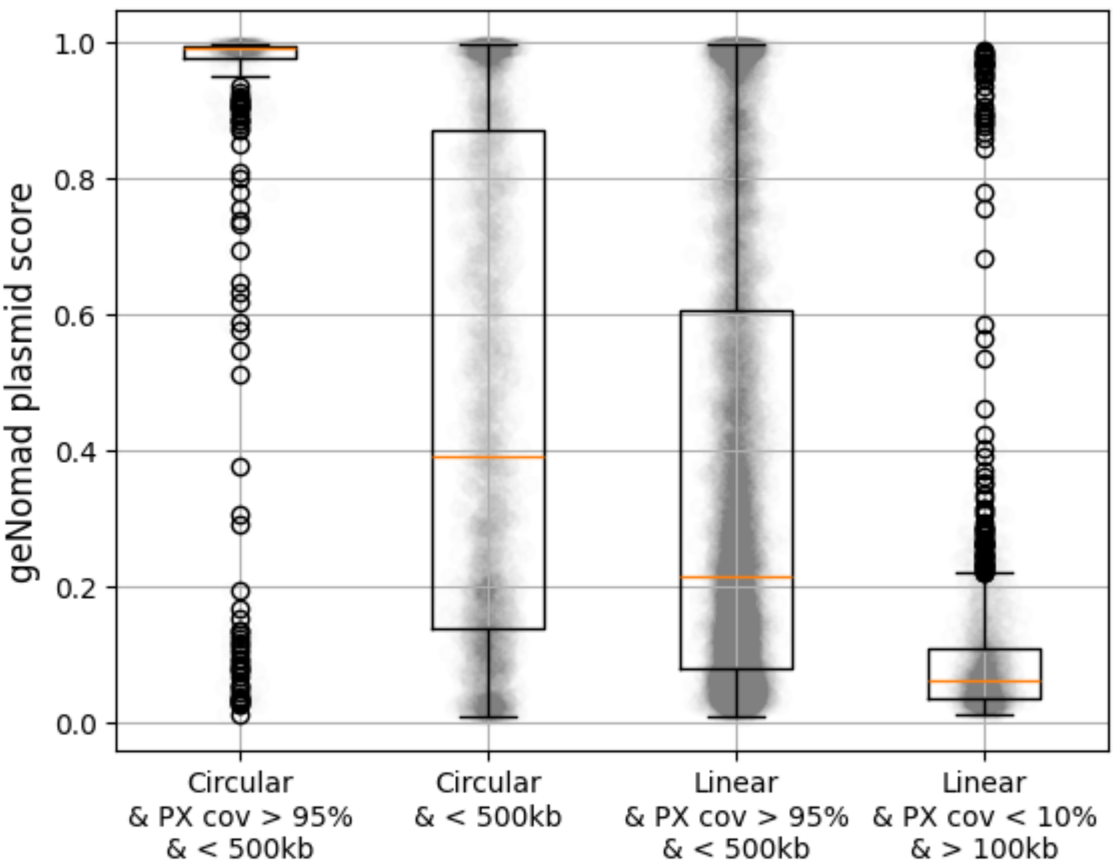


**Supplementary Figure 20. GeNomad plasmid scores against decreasing plasmid evidence**. (left) long-read circular contigs at least 95% covered by metaplasmidomics reads, and shorter than 500kb, (left-center) long-read circular contigs shorter than 500kb, (right-center) long-read linear contigs at least 95% covered by metaplasmidomics reads, and shorter than 500kb, (right) long-read linear contigs at maximum 10% covered by metaplasmidomics reads, and longer than 100kb. PX cov: plasmidomic reads coverage.

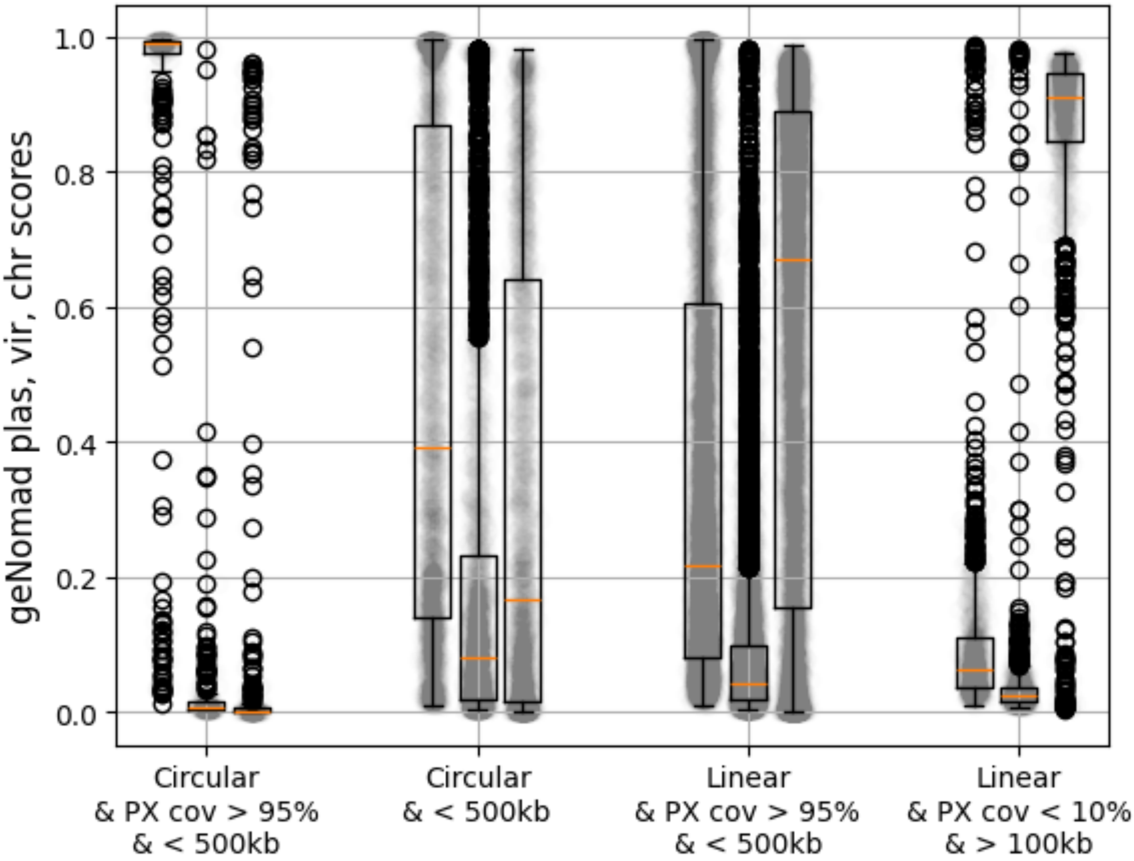


**Supplementary Figure 21. GeNomad plasmid, virus, and chromosome scores against decreasing plasmid evidence**. (left) Long-read circular contigs at least 95% covered by metaplasmidomics reads, and shorter than 500kb, (left-center) long-read circular contigs shorter than 500kb, (right-center) long-read linear contigs at least 95% covered by metaplasmidomics reads, and shorter than 500kb, (right) long-read linear contigs at maximum 10% covered by metaplasmidomics reads, and longer than 100kb. PX cov: plasmidomic reads coverage. For each plasmid evidence group, the boxplots show the distribution of plasmid, virus, and chromosome geNomad scores from left to right.
